# Identification of heterotic group-specific haplotypes and impact of residual inbreeding on grain yield of maize elite hybrids

**DOI:** 10.64898/2026.06.15.732226

**Authors:** Romain Kadoumi, Nicolas Heslot, Fabienne Henriot, Alain Murigneux, Mathilde Berton, Laurence Moreau, Alain Charcosset

## Abstract

Modern hybrid maize (*Zea mays L.*) breeding programs are based on the management of distinct complementary heterotic groups to maximize heterosis in high-performing hybrids. This practice lowers shared genetic segments and increases divergence between groups to limit inbreeding in hybrids. However, most breeding programs have not always enforced strict separation between heterotic groups in the past. Competitor commercial hybrids were notably a common elite germplasm source for inbred development, which would diminish divergence between groups.

This study proposes a new haplotype-based approach to assess hybrids’ residual inbreeding based on parental similarity. The new haplotype method has a stronger significant negative effect on hybrids’ grain yield than raw SNP data. Evaluation of modern experimental hybrids uncovered related inbreds contributing to superior rates of residual inbreeding. Analysis of these inbreds revealed haplotype transfers between heterotic groups, originating notably from the use of a Stiff Stalk-Iodent commercial hybrid as breeding starts material in both Stiff Stalk and Non-Stiff Stalk breeding populations. The introduction of this intergroup parent generated heterotic-group-specific haplotype migration between crossing pools. These fragments caused significant genome-wide residual inbreeding in experimental hybrids across selection cycles.

This study highlights the necessity for accurate evaluation of external sources of diversity to minimize haplotype transfers and admixture between crossing pools. We demonstrate the consequences of using commercial hybrids in inbred development, particularly regarding residual inbreeding, and their effects on hybrid performance. Insights from these results can assist breeders in optimizing the choice of parents for introducing genetic diversity in a reciprocal recurrent selection scheme.

**KEY MESSAGE:** Haplotype-based hybrid’s parental similarity better predicts grain yield than marker-based identity-by-state. Utilization of commercial hybrids as breeding start material resulted in higher hybrid residual inbreeding even after several selection cycles

## INTRODUCTION

The invention of hybrids (Shull 1908) and early observations of hybrid vigor by East (1907) and Shull (1909) positioned maize (Zea mays subsp. mays) as a model crop for studying heterosis and implementing novel hybrid selection strategies. Hybrid breeding relies on the controlled crossing of genetically distinct inbred lines to produce high-performing hybrids. Since the 1920s, hybrids have exhibited superior agronomic traits, including higher yields, greater uniformity, and reduced susceptibility to environmental variability, when compared to traditional landraces and open-pollinated varieties (Crow 1998; Troyer 2006). Often opposed to heterosis, inbreeding is a critical consideration in maize hybrid breeding. Inbreeding is the reduction of biological fitness and vigor resulting from the mating of genetically related individuals (or self-reproduction). It arises from the accumulation of deleterious recessive alleles in a population (Charlesworth and Willis 2009). When inbreeding depression is important, it is usually managed in hybrid breeding crops by creating distinct pools of individuals (so called heterotic groups in maize) to maximize genetic divergence between them. This concept parallels animal breeding practices, such as terminal crosses used in poultry and pork production, which also exploit hybrid vigor to enhance productivity and uniformity, and avoid the harsh consequences of animal inbreeding depression (Lasley 1977; Lo et al. 1997; Vanvanhossou et al. 2025). In hybrid populations, residual inbreeding refers to the inadvertent retention of homozygosity in terminal hybrids, resulting from the cross of incompletely divergent inbred lines. This phenomenon can thus significantly attenuate the heterotic advantage by limiting genetic complementation.

To persistently exploit heterosis and limit hybrid residual inbreeding, breeding schemes have been adapted to systematically favor the crossing of parental inbreds with divergent complementary genetic backgrounds. This tendency prompted the interest in tester-based reciprocal recurrent selection schemes (RSS). It was rapidly accepted by both private-commercial and public programs and contributed to their growing dynamics (Hallauer et al. 1983; Gracen 1986). This approach, first conceptualized by Hull (1945), involves the cyclical and simultaneous selection of two genetically divergent elite inbred groups. Each group serves as source material for within-group selection (i.e., inbred recycling) and as testers to evaluate intergroup hybrid crosses capabilities with the opposite divergent population of the crossing pattern. This process enables the stacking of favorable divergent alleles, improving performances (e.g., yield, adaptative traits, etc.) (Comstock et al. 1949; Menz Rademacher et al. 1999; Hinze et al. 2005; Edwards 2011). At a population level, this scheme creates a heterotic population structure, with increasing genetic differentiation over time (Melchinger et al. 1991; Reif et al. 2005).

While RSS undoubtedly contributed to increased performances and sustained genetic gain (Penny and Eberhart 1971; Crow 1998), the continuous recycling of highly related elite inbreds within groups has been shown to narrow effective genetic diversity, cause genetic variability loss, and limit genetic gain in breeding schemes (McCouch et al. 2013; Mickelbart et al. 2015; Allier et al. 2019). Strategies based on the integration and monitoring of novel genetic resources were proposed to mitigate this issue. It notably involves the use of public lines, landraces, expired Plant Variety Protection (ex-PVP) lines, tropical and exotic material (Guo et al. 2021; Smith et al. 2022; Sanchez et al. 2024). However, in the context of hybrid schemes, these approaches may also negatively impact heterotic pattern divergence and the heterotic group structure, if related source materials are introduced into complementary heterotic groups (Charlesworth 2003; Verhoeven et al. 2011). In maize breeding, commercial F1 hybrids represent alternative genetic resources for inbred breeding schemes. Historically, in authorized geographical regions, this method enabled breeders to access competitors’ commercial elite germplasm through selfing and recombinations. For instance, during the 1980-2000 period, commercial hybrids were identified in the pedigree of 3% of American ex-PVP lines (Mikel and Dudley 2006). According to Mikel (2011), the most notable example is the introgression of the novel Iodent germplasm within Monsanto breeding programs via the Dekalb DK3IIH6 line, a major genetic contributor derived from a commercial hybrid, according to its PVP certificate (PVP9300087). However, this approach remains relatively rare due to concerns about disrupting heterotic group differentiation and the increasing germplasm logistical complexity of managing hybrid-derived admixed inbreds (e.g., risks of heterotic pools admixture). Bernardo (2001)’s study on the incorporation of interheterotic group crosses for inbred development and Zhang et al. (2024) evaluation of interheterotic group composites highlighted these challenges, notably the search for suitable testers.

In recent years, the advances in sequencing technologies and analytical frameworks have enabled the use of haplotypic information in quantitative genetics, as an alternative to the use of individual loci. The incorporation of multi-loci data has improved the understanding of complex traits and offered new insights into population structure, genetic events (e.g., mutation, selection), linkage patterns and functional diversity for plant breeding (Lorenz et al. 2010; Romay et al. 2013; Mayer et al. 2020). A haplotype block defines a genomic region that comprises a set of closely linked phased alleles that tend to be inherited together with a small probability of recombination. A haplotype allele is a combination of phased SNP alleles found in a specific haplotype block, which are in high linkage disequilibrium (LD) (Daly et al. 2001; Gabriel et al. 2002; Clark 2004). Compared with individual SNP markers, which capture identity-by-state (IBS), haplotypes are expected to be more informative on the finer-scale recombination history and ancestral identity-by-descent (IBD), providing a more accurate representation of genetic relatedness between individuals (Kim and Yoo 2016). Therefore, applied to commercial plant breeding, haplotyping enables tracking chromosomal fragments shared between individuals. This is especially relevant in modern programs to follow ancestral genomic region conservation, historical introgressions and selective sweeps found in elite parental inbreds (Coffman et al. 2020; Bornowski et al. 2021).

In this study, we first introduce a novel haplotype-based metric to quantify hybrids’ residual inbreeding by analyzing parental genetic similarities. This method is applied to a dataset of experimental hybrids, evaluated in 2024, from a commercial European breeding program. It was used to investigate the relationship between residual inbreeding and hybrid performances. From these results, we then identified the ancestral haplotypic origin leading to increased rates of residual inbreeding in a specific inbred family, obtained from the use of a commercial hybrid. We aim to (1) quantify shared genetic fragments between the major heterotic groups in the dent maize crossing pattern (Stiff Stalk x Non-Stiff Stalk); (2) trace the origins and evolutionary trajectories of these shared haplotypes; and (3) assess the impact of common genetic fragments on residual inbreeding and its consequences on hybrid performances. The article provides a unique proof-of-concept for the use of haplotype-based indicators to detect and quantify residual inbreeding in hybrids, as well as to predict its consequences on maize grain yield. This research offers insights for improving modern hybrid breeding programs, through the optimization of inbred development and hybrid crosses.

## MATERIAL AND METHODS

### Plant material, pedigree extraction and used datasets

To evaluate the extent of hybrids’ parental similarity in modern elite maize hybrids and assess its impact on grain yield, a primary dataset (D1) was assembled consisting of 2,390 experimental hybrids, evaluated in Europe during the summer of 2024 (Fig. 1). The hybrids follow the Stiff Stalk (SS) x Non-Stiff Stalk (NSS) heterotic pattern and were obtained from the cross of 332 SS with 381 NSS parental inbred lines (Supplementary Fig. 1). NSS results from the recent merging of Iodent (IDT) and Lancaster heterotic groups (Kadoumi et al. 2025). These hybrids were tested for the European mid-early/mid-late breeding program managed by *Limagrain Field Seeds*, which focuses on high-yielding grain dent varieties, adapted for southern France, Central and Eastern Europe. From this initial set, we observed that a number of hybrids showing high residual inbreeding had parents that shared one commercial hybrid common ancestor. We then identified all hybrids, whose two parents share this common ancestor in their pedigrees. This constitutes a subset of 97 hybrids (D1S) with variable residual inbreeding level, obtained from the crosses between 53 related inbreds (25 SS and 38 NSS parents) (Fig. 1; Supplementary Fig. 2).

**Fig. 1.**
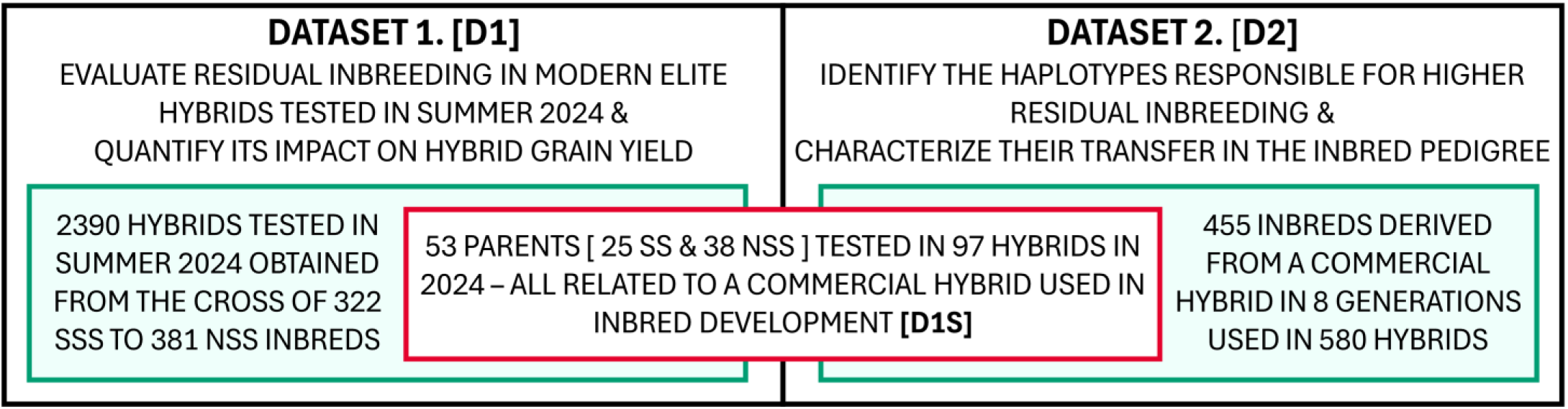
Synthetic representation of the datasets (D1 and D2) used in this study. The 53 parents used in the 97 hybrids are the common material between both datasets (D1S)

To investigate the genetic background contributing to the elevated inbreeding values in the subset of 97 hybrids, a second dataset (D2) was constructed starting from the most recent common ancestor between the two heterotic groups parental inbreds: a competitor commercial hybrid. This distinctive parental material, hereafter noted COMPHYB, was commercialized in France in the 1980s’ and is putatively a Stiff Stalk type-B37 x Iodent hybrid. We then extracted the complete inbred progeny of this “founder” individual, which contributed to the development of inbreds in both Stiff Stalk and Iodent heterotic groups. This dataset allowed for a more detailed exploration of ancestral contributions and potential transfers among inbreds of different heterotic groups.

On the SS side, COMPHYB was crossed to two SS inbreds (i.e., a Limagrain B73-type and the public line A632), creating two “breeding topcrosses” (noted TC1 and TC2). COMPHYB was also selfed 4 times to derive inbred lines (noted SELF1 to SELF4). These selfed inbreds were later used as parents in other inbred breeding pipelines, including the Iodent inbred pipeline (Supplementary Fig. 3; Supplementary Table 1). Consequently, the second dataset consists of a total of 455 inbreds derived from COMPHYB over 7 inbred breeding cycles, which led to 580 tested hybrids, including the 97 hybrids tested in 2024 (present in the first dataset D1S). To better characterize the material obtained from COMPHYB, 7 reference inbreds were added for comparison, including the two TC SS parents, 4 SS founder lines issued from the original BSSS population (B14, B37, B73, and B64), and an Iodent group check MBS847. To characterize the evolution of COMPHYB material in the progeny, inbred lines were agglomerated according to the number of inbred selection cycles realized after the first cross with COMPHYB in C0 (Supplementary Fig. 3). To evaluate the impact of breeding and selection cycles on the evolution of residual inbreeding in hybrids, we calculate the complete pairwise comparisons between the 455 inbreds, leading to 106,491 potential pairwise single-cross hybrids, including both intragroup and intergroup hybrids (D2HP). We compare these hypothetical crosses to the 580 tested intergroup hybrids (D2HG) and the subset of tested hybrids in 2024 (D2HM).

### Genotyping, marker processing and genome segmentation into haplotypes

Genotyping was performed for the inbred lines of both datasets using the *Limagrain*-designed Affymetrix genotyping array, which consists in 18,480 single nucleotide polymorphisms (SNP) markers, selected to represent polymorphism in temperate European germplasm (Kadoumi et al. 2025). Missing data were imputed for each dataset and genotypes were phased to manage residual heterozygotes during haplotyping, using the software BEAGLE v5.4, with default settings (Browning et al. 2018). No pruning for minor allele frequency nor marker heterozygosity was applied. The first dataset resulted in 713 inbreds (332 SS and 381 NSS parents) genotyped across 18,439 SNP markers. The second dataset obtained comprised 462 inbreds (206 SS, 56 admixed inbreds (ADMX) and 200 IDT) analyzed across 18,439 markers.

Haplotype blocks were defined for each dataset D1 and D2, using jointly all inbreds (SS and NSS) without considering population structure. Chromosome segmentation in consecutive haplotype blocks was realized using the software *HaploView* 4.2, with the solid spine algorithm (Barrett et al. 2005). Compared to the other proposed methods (e.g., “Gabriel confidence interval” (Gabriel et al. 2002), “Four-Gametes Test” (Wang et al. 2002)), this algorithm attempts to maximize block length by only considering the LD of the spine (i.e., the outermost correlation) of the haplotype block. The minimum pairwise marker correlation threshold was D’= 0.2 to maximize block length and create common haplotype blocks between the two crossing pools (Mayer et al. 2017). The maximum pairwise distance threshold was set to 50 Mb due to limited computation capabilities. No marker pruning based on allele frequency nor linkage disequilibrium was realized, and all 18,439 SNPs were assigned into haplotype blocks. Single-SNP haplotypes were kept.

### Residual inbreeding assessment

Residual inbreeding (RIB) of a hybrid (intergroup or intragroup) was calculated at a local and global scale. At a local genomic scale, for each haplotype block, we calculated the probability of each parental haplotype matching the other parent (i.e., probabilistic Hamming distance (Hamming 1980)). To manage residual heterozygosity in non-doubled haploid inbreds, we assumed that both haplotypes present in a parent have an equal probability of transfer to its hybrid progeny. Hence, up to four potential unique hybrid profiles can exist per pair of parents. We assumed that the 2 possible phased haplotypes per parent segregate in the hybrid without any recombination. To scale up RIB at a global genome level, we summed each haplotype block probability of similarity over the genome. We controlled for haplotype block size by weighting each block by its number of SNP markers and normalizing by the total number of markers. The indicator is thus a percentage of similarity between parental pairs. We also calculated for each pair of parents the biallelic SNP-based Identity-by-State (IBS) similarity metric. IBS is computed as the number of matching homozygote SNPs and half the number of matching heterozygotes SNPs, divided by the total number of markers (i.e., here 18,439 SNPs).

### Phenotypic data and Adjusted means in dataset D1

We used the dataset D1 phenotypic data to evaluate the impact of residual inbreeding on hybrid grain yield. The 2,390 hybrids were assessed in multiple trials across Europe in the summer of 2024 for grain yield at 15% grain moisture (in quintals per hectare). We filtered out observations of hybrid crosses present in a single location. Locations with strictly less than 10 unique hybrid crosses were discarded. The resulting dataset was thus composed of 31,994 observations in 208 locations, including 1,357 for the subset of 97 hybrids with potential residual inbreeding from COMPHYB. As the retrieved data were already adjusted for intra-field effects, adjusted means (also called least-square means or ls-means) of the hybrids were computed for the grain yield to correct location-dependent environmental effects. The following statistical model was used considering the hybrid genetic effect as fixed and the location as random:

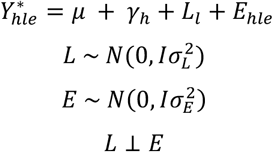

Where 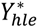 is the phenotype of the hybrid *h* in the location *l* and replication *e, μ* is the intercept, *γ*_ℎ_ is the fixed effect of the hybrid *h, L*_*l*_ is the random effect of the location l with 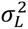 variance and *E*_ℎ*le*_ is the random error of the model with 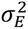 variance.

### Variance decomposition, fixed effects significance and kinship matrices computation

Variance decomposition was realized to gauge the proportion of parental genetic variance components in the first hybrids dataset (D1), following the GCA-model of González-Diéguez et al. (2021). Four models were tested: the A model (i.e., only parental additive effects), the A(AA) model (i.e., A model with hybrid additive x additive epistatic effect), the AD model (i.e., A model with hybrid dominance effect) and the AD(AA) model (i.e., A model with both hybrid additive-additive epistatic and dominance effects). To estimate the significance of parental similarity in genomic mixed models, and its consequences on variance components, we implement different models including or not IBS or RIB similarities as fixed covariates; M0: null model/no fixed effects, M1: IBS, M2: RIB, M3: IBS and RIB. The following model is the most complete / saturated AD(AA) with both fixed effects, and was applied to the hybrid grain yield adjusted mean values:

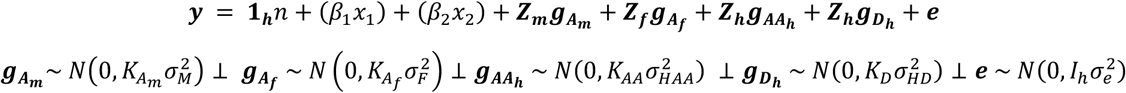

Where ***y*** is the vector of adjusted grain yield means of the h phenotyped hybrids, **1**_***h***_*n* is the vector intercept of size *h*, ***g***_***Am***_ (***g***_***Af***_) is the vector of random additive male (female) GCA effects, and ***g***_***AAh***_ and ***g***_***Dh***_ are the random hybrid epistatic additive-additive, and dominance SCA effect, respectively. *β*_1_ and *β*_2_are the fixed linear beta regression effects on the *x*_1_ (IBS) and *x*_2_ (RIB) covariates corresponding to the RIB or IBS similarity coefficient between the two parental lines of the hybrid. ***e*** is the vector of error terms of 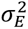 variance. ***Z***_***m***_, ***Z***_***f***_, and ***Z***_***h***_ are the incidence matrices that relate the phenotypic records to the genetic effects. Kinship matrices for the parental additive GCA effects (*K*_*Af*_ and *K*_*Am*_) were computed following the Genomic Relationship Matrix (GRM) method 1 from VanRaden (2008). The hybrid epistatic additive-additive kinship matrix (*K*_*AA*_) was produced from the crossing of the respective parental coefficient (*ij* and *i’j’*)of the hybrids *h* and *h’* (i.e., Hadamard product) as follows (Henderson 1985; Technow et al. 2014), and divided by the mean value of the matrix (i.e., trace divided by the number of hybrids). The hybrid dominance SCA kinship matrix (*K*_*SCA*_) was computed using the formula from González-Diéguez et al. (2021), which uses the parental alleles frequencies. To ensure that all kinships were positive definite, an additional 10^-6^ was added to the diagonal. To gauge the significance of the fixed effects, ANOVA Wald tests were performed (with a risk level α = 0.05) using the R function *AnovaTest()* of the package *MM4LMM*. When two fixed effects are tested, a conditional test is implemented. All models were computed using the function *MMEst()* of the R package *MM4LMM* (version 3.0.3), which performs a “restricted Maximum Likelihood” (ReML) inference of variance components in mixed model, using a Min-Max algorithm (Laporte et al. 2022; Laporte et al. 2024). To compare the different models with identical fixed effects, consecutive pairwise Likelihood Ratio Test and differences in AIC were computed.

### Population Structure of the second dataset

Principal Component Analysis (PCA) was used to identify the largest sources of variation and to visualize the population structure of the second dataset (D2). PCA was run with the R package smartsnp (version 1.1.0) (Herrando-Perez et al. 2021) and the function smart_pca(), which utilizes the eigenstrat method (Price et al. 2006). The top 3 principal component axes were retrieved. Inbreds assignation to specific genetic groups was realized using the Bayesian clustering software *ADMIXTURE* (version 1.3.0) (Alexander et al. 2009). The analysis was conducted with default parameters and 10 cross-validations for unsupervised clustering at K (number of clusters) values equal to 3, to mimic SS, IDT, and the potential intergroup SS-IDT admixed inbreds. As the marker set was considered representative of the analyzed diversity, no marker was removed due to allele frequencies, heterozygosity, or linkage disequilibrium.

### Haplotype sharing, origin assignation and expected residual inbreeding in hybrids

For the second dataset D2, as COMPHYB parents are unknown, we reconstructed the COMPHYB hybrid profile through its progeny. In this dataset, genetic material from COMPHYB only exists through its six first-cycle inbreds (4 selfings and 2 breeding topcrosses). We estimated the share of COMPHYB material as the sum of 1) all possible haplotypes derived from selfings and 2) all haplotypes unique to the TC inbred and different from their SS parents. To find the potential origin of haplotype transfers between heterotic groups, each haplotype was assigned to a genetic group based on the earliest inbred carrying it. The reference inbreds’ genealogy was retrieved using their release year found in the USDA GRIN-Global National Plant Germplasm System database. To evaluate the potential impact of material transfer between and inside genetic groups and gauge the impact of selection decisions, we calculated the parental similarities (IBS and RIB) for all pairwise combinations of the dataset, representing 106,491 potential single-cross hybrids (intragroup and intergroup, noted D2HP), of which 580 were tested between 2008 and 2024 at different selection stages (D2HG). Experimental hybrids tested in 2024 (D2HM) are the same as the highlighted hybrids found in the dataset 1 (D1S)

In the precedent method sections, unless stated otherwise, all analyses were done in R 4.4.1 (R Core Team 2024), and visualizations were plotted using the R package ggplot2 (version 3.5.1) (Wickham et al. 2024).

## RESULTS

### Hybrids’ parental similarity in dataset D1

Haplotype-based Residual Inbreeding (RIB) of these hybrids was evaluated using the weighted average similarity between parental inbreds, based on a set of 1326 LD-defined haplotype blocks. The average size of the haplotype blocks was 14 consecutive markers (standard deviation = 12 SNPs). RIB was comprised between 0.024 and 0.178, with an average of 0.054 ± 0.017. Average Identity-by-State (IBS) between parents of tested hybrids for dataset D1 was 0.581 ± 0.011 (min = 0.553; max = 0.641), representing on average 10,713 common SNP markers between hybrid parents. Analysis of both parental similarity indicators showed that IBS was normally distributed, while RIB was left-skewed with an over-representation of low RIB values (Supplementary Fig. 4). Pearson correlation between IBS and RIB, for the entire dataset, was 0.88 (p-value < 2.2E-16), and 0.91 (p-value < 2.2E-16) when focusing on the subset of 97 hybrids (Fig. 2). Comparison of parental similarity metrics revealed significantly superior values for both IBS (global = 0.580; subset = 0.587; p-value = 9.5E-09) and RIB (global = 0.054; subset = 0.066; p-value = 2.0E-07) for the subset of 97 hybrids in comparison to the global dataset. Application of a linear model to explain RIB using only IBS as a fixed eff ect indicated a positive regression coefficient of 1.37 and an intercept of -0.742.

**Fig. 2.**
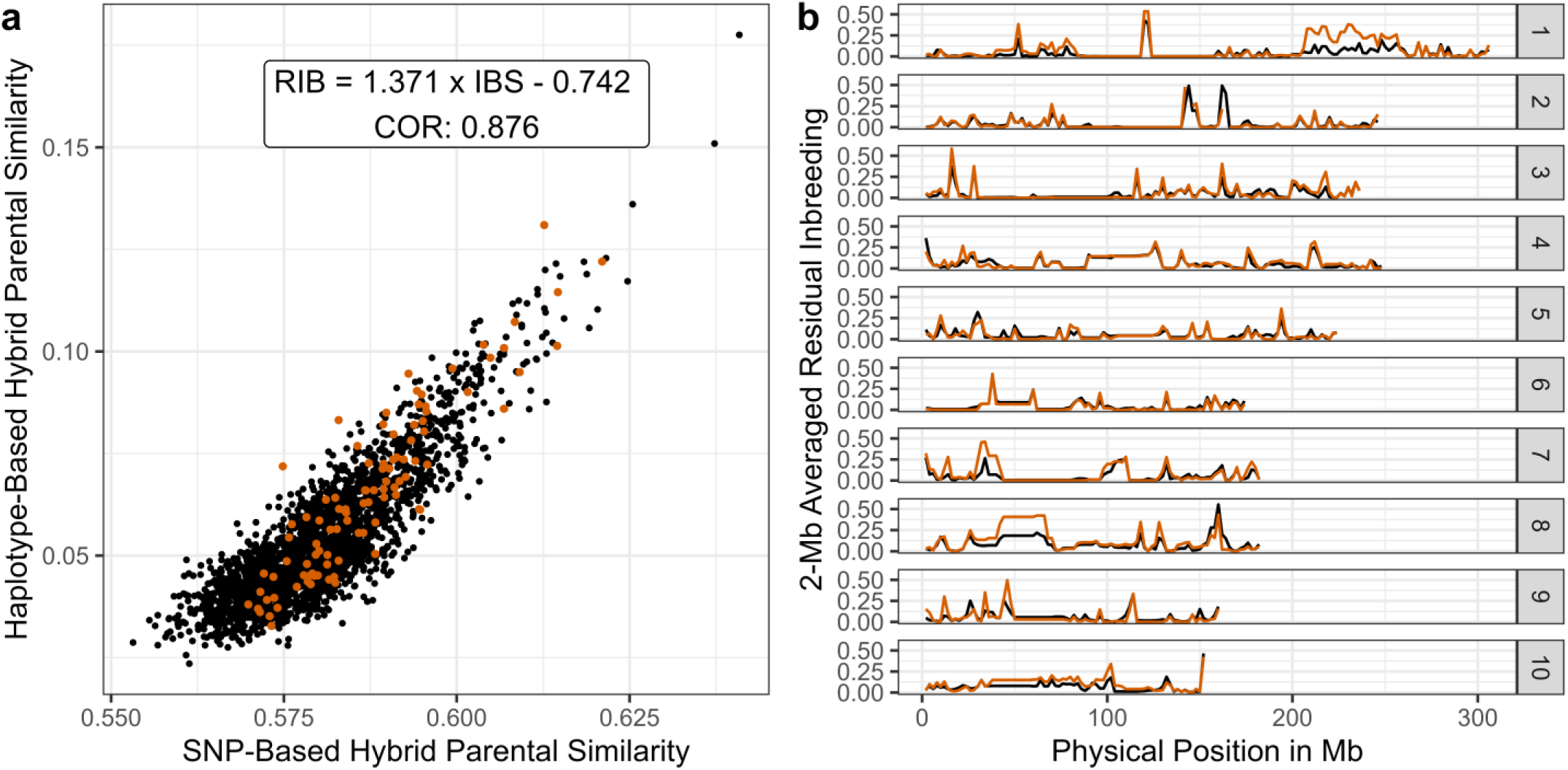
Overview of 2024 experimental hybrids’ residual inbreeding. a) Scatter plot between IBS and RIB. b) Evolution of RIB along the genome Color scheme separates the global 2024 experimental hybrids dataset D1 (black) to the subset of 97 hybrids (D1S), where both parents of the 97 hybrids originate from a common ancestor (orange)

Genome-wide mapping of local hybrids’ residual inbreeding (RIB) was monitored to observe potential outlier regions with superior parental similarity and highlight the differences between the global 2024 experimental hybrids and the subset of 97 hybrids (Fig. 2). This approach highlighted 3 chromosomic sections in the 97 hybrids, with consecutively multi-haplotype block high residual inbreeding: on chromosome 1 telomere at 225 Mb, on chromosome 4 centromere at 100 Mb, which can also be found in the complete panel of 2024 hybrids, and on chromosome 8 at 50 Mb. The chromosome 8 region had the highest average residual inbreeding, with 42.3% of the 97 hybrids (i.e., 41 hybrids) having homozygous haplotypes.

### Impact of residual inbreeding on performance and genomic prediction

Computation of grain yield adjusted means showed an average of 125 qt/ha (quintals per hectare), with a standard deviation of 6.3 qt/ha. Location random effect standard deviation was 32.35 qt/ha. As a preliminary step and outside of genomic mixed models fitting, we analyzed the impact of parental similarity on hybrid grain yield using the Pearsons’ correlation test. It demonstrated weak negative correlation (Supplementary Fig. 5). Decomposition of the genetic variance into its parental GCA and interaction components (epistasis and SCA), indifferent of the considered model and fixed effect part, revealed a predominance of female GCA (i.e., SS parent) variance in the total genetic variance (Table 2). In the most complete null model AD(AA), female additive GCA variance reached 51.7%, while SCA interaction genetic variance, measured by both epistatic additive-additive and dominance effects, was 32.5%. Across all models’ fixed effect types (null, IBS, RIB, IBS+RIB), complexification of the statistical models to incorporate SCA interaction effects highlighted a systematic decrease in the models’ error, improved the models’ loglikelihood and had a significant pairwise LRT at the 5% p-value threshold. Insertion of a single fixed parental similarity effect (IBS or RIB) was significant at 5% p-value with an ANOVA Type-3 Wald test, and had a delta AIC above 10, compared to the null model (Supplementary Table 2). Beta effect values were strongly negative for grain yield, with an average of -50.84 for IBS and -32.71 for RIB. This represents a loss of around 1 qt/ha for every 2% of increased parental similarity for IBS. While having non-significant p-values according to the Wald test, the combination of both fixed effects (IBS and RIB) was supported with partial evidence, with a superior difference in AIC compared to models including only IBS or RIB. The summary results of the models can be found in Table 2.

**Table 1.** Consequences of the incorporation of fixed parental similarity effects on genomic evaluation models. Summary results for the sixteen models and variance decomposition for the hybrid grain yield adjusted means. Variance components are decomposed between the parental female additive GCA variance (A-F), the parental male additive GCA variance (A-M), the hybrid epistatic additive-additive variance (AA), the hybrid dominance SCA variance (D) and the error variance (E). For each fixed effect (FE) applied, four mixed models were used: a model with only parental additive GCA variance (A), the parental additive GCA and dominance SCA model (AD), the parental additive GCA and epistatic additive-additive model (A(AA)) and the complete model (AD(AA)) with the parental additive GCA, the dominance SCA and the interaction additive-additive variances.

| FE | Model | LL | Variance Components |  |  |  |  | IBS |  | RIB |  |
| --- | --- | --- | --- | --- | --- | --- | --- | --- | --- | --- | --- |
|  |  |  | A-F | A-M | AA | D | E | Beta | P-value | Beta | P-value |
| Null | A | -5008.748 | 12.92 | 5.21 | - | - | 17.88 | - | - | - | - |
|  | AD | -4982.890 | 11.31 | 2.85 | - | 4.26 | 14.66 | - | - | - | - |
|  | A(AA) | -4977.555 | 10.87 | 4.07 | 6.67 | - | 13.26 | - | - | - | - |
|  | AD(AA) | -4975.090 | 10.65 | 3.27 | 4.73 | 1.96 | 13.15 | - | - | - | - |
| IBS | A | -5000.831 | 12.49 | 5.35 | - | - | 17.81 | -48.090 | 1.2E-03 | - | - |
|  | AD | -4976.133 | 10.97 | 2.98 | - | 4.22 | 14.64 | -54.554 | 5.8E-03 | - | - |
|  | A(AA) | -4970.810 | 10.47 | 4.22 | 6.64 | - | 13.24 | -48.822 | 5.0E-03 | - | - |
|  | AD(AA) | -4968.399 | 10.28 | 3.43 | 4.78 | 1.89 | 13.13 | -51.899 | 5.9E-03 | - | - |
| RIB | A | -4999.886 | 12.59 | 5.34 | - | - | 17.78 | - | - | -32.502 | 2.4E-04 |
|  | AD | -4975.875 | 11.14 | 3.00 | - | 4.16 | 14.64 | - | - | -35.086 | 2.4E-03 |
|  | A(AA) | -4970.894 | 10.65 | 4.23 | 6.48 | - | 13.30 | - | - | -30.644 | 3.1E-03 |
|  | AD(AA) | -4968.477 | 10.46 | 3.42 | 4.60 | 1.92 | 13.18 | - | - | -32.600 | 3.5E-03 |
| IBS + RIB | A | -4996.507 | 12.57 | 5.35 | - | - | 17.79 | -5.550 | 8.5E-01 | -29.659 | 8.6E-02 |
|  | AD | -4972.22 | 11.08 | 3.00 | - | 4.19 | 14.64 | -16.036 | 6.5E-01 | -27.378 | 1.8E-01 |
|  | A(AA) | -4967.247 | 10.57 | 4.24 | 6.54 | - | 13.28 | -18.474 | 5.7E-01 | -21.313 | 2.8E-01 |
|  | AD(AA) | -4964.753 | 10.38 | 3.43 | 4.67 | 1.92 | 13.15 | -19.530 | 5.7E-01 | -22.959 | 2.6E-01 |

\* FE: Fixed Effect / LL: LogLikelihood

**Table 2.**
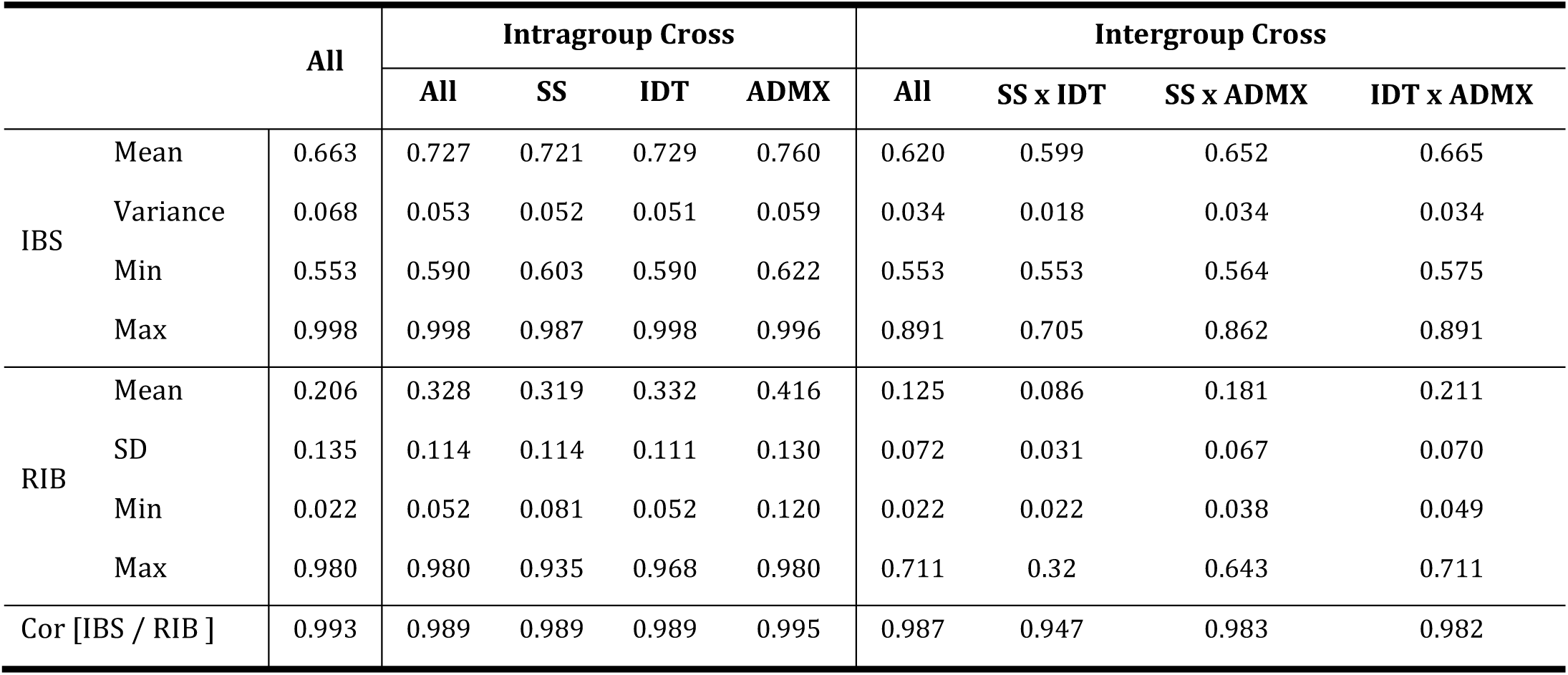
Parental similarity summary metrics of intragroup and intergroup crosses from the complete pairwise combinations of all inbreds.

### Population structure, haplotypic dataset and pedigree relationships in dataset D2

Principal Components Analysis of the second dataset genotyping matrix revealed three linked clusters corresponding to Stiff Stalk (SS), Iodent (IDT), and an Admixed (ADMX) group, with inbreds connecting SS to ADMX and ADMX to IDT (Fig. 3). Proportion of variance explained by the top three first PCs was, 13.88%, 4.04%, and 3.31% respectively, reaching a total of 21.23%, with PC1 separating the global dataset into SS – ADMX – IDT in a gradient manner, PC2 splitting ”pure materials” from admixed ones, and PC3 segmenting the population according to SS precocity. Haplotype block construction revealed 1232 haplotypes blocks, with an average size of 15 consecutive SNPs (SD: 13 SNPs; min: 2 SNPs; max: 116 SNPs). On average, there were 25.6 haplotypes alleles per haplotype block (SD: 16 haplotypes; min: 2 haplotypes; max: 133 haplotypes), representing a total of 31,587 haplotypes in the total dataset.

**Fig. 3.**
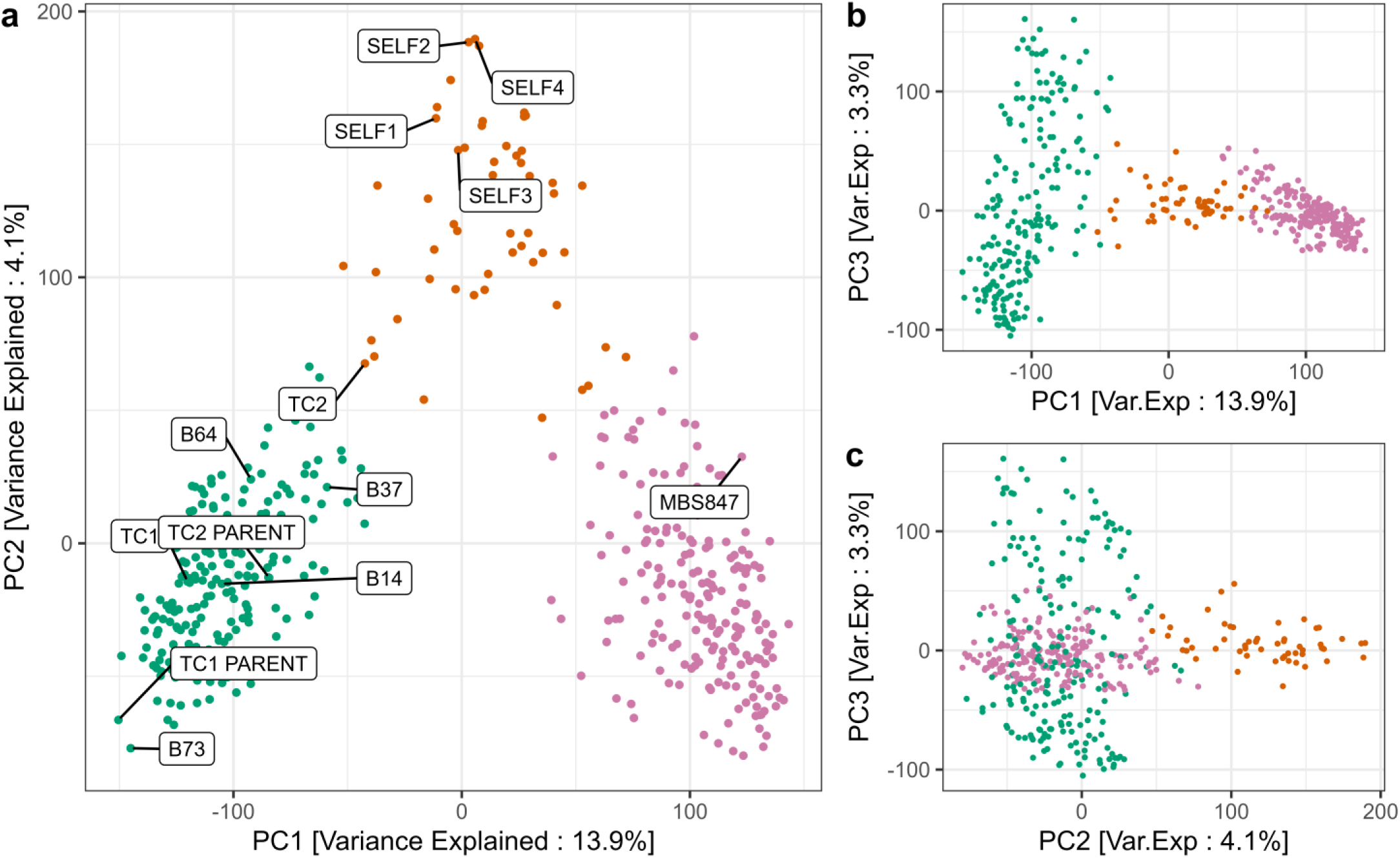
Evolution of the germplasm’s population structure (for the inbred dataset D2) visualized by SNP-based PCA scatterplots (PC1 and PC2). Color scheme represents the ADMIXTURE assigned genetic group: Stiff Stalk (green), Iodent (pink) and Admixed (orange)

Classification of the dataset’s pedigree architecture highlighted unbalanced distribution of inbreds per cross type (SELF or TC), with 326 inbreds derived from the selfing of the hybrid, and 141 inbreds obtained from the breeding TCs. Twelve inbreds, assigned to the SS, and obtained between C3 and C7, have pedigree links to both SELF and TC C0 starting crosses. Genetic group assignation using ADMIXTURE revealed that 44.0%, 43.7% and 12.3% of lines of D2 were assigned to SS, IDT and ADMX, respectively. Finer decomposition of pedigree relationships reveals that 85.7 % of all inbreds trace back to only 2 out of the 6 C0 inbreds: SELF2 and TC1.

### Haplotype sharing between founders, the commercial hybrid and its progeny

To characterize haplotype transfers between groups and notably those corresponding to fragments originating from COMPHYB, we analyzed haplotype sharing and its evolution between the group founder inbreds, COMPHYB and its progeny. As COMPHYB genotype is unknown, we deciphered it through a haplotype analysis of the first-cycle progeny (C0). For TC derived materials, the contribution of COMPHYB was obtained from the TCs haplotypes not assigned to their respective SS parents (LG B73-type and A632). It was equal to 32.9% and 58.7% for TC1 and TC2, respectively. For TC1, the proportion of group reference haplotypes, was 80.8% for BSSS, and 14.7% for MBS847. For TC2, the proportion of group reference haplotypes, was 57.0% for BSSS and 20.6% for MBS847 haplotypes. For SELF, which were 100% COMPHYB-derived, the proportion of group reference haplotypes reached on average 44% for BSSS (i.e., on average 23.3% of B64, 23.2% of B37, 15.5% for B14 and 8.0% for B73) and 27.2% for MBS847. Hence, around 28.8% of SELF haplotypes were not explained by the group reference material and were considered as corresponding to the COMPHYB haplotypes.

The evolution of the proportion of COMPHYB haplotypes was monitored across its different progeny selection cycles (Fig. 4). Across all selection cycles, the average frequency of COMPHYB haplotypes was 35.4%, with a standard deviation of 15%. We observed a clear decrease in the frequency of COMPHYB haplotypes across selection cycles, starting at an average of 66.7% (42.9% in the SS; 78.6% in the ADMX) at C0-1 and reaching a plateau at 28.1% at C6-7. Contrary to COMPHYB haplotypes, those issued from group reference lines (B14, B37, B73 and B64 for BSSS and the Iodent check) remained relatively stable in their respective genetic groups (Supplementary Table 3).

**Fig. 4.**
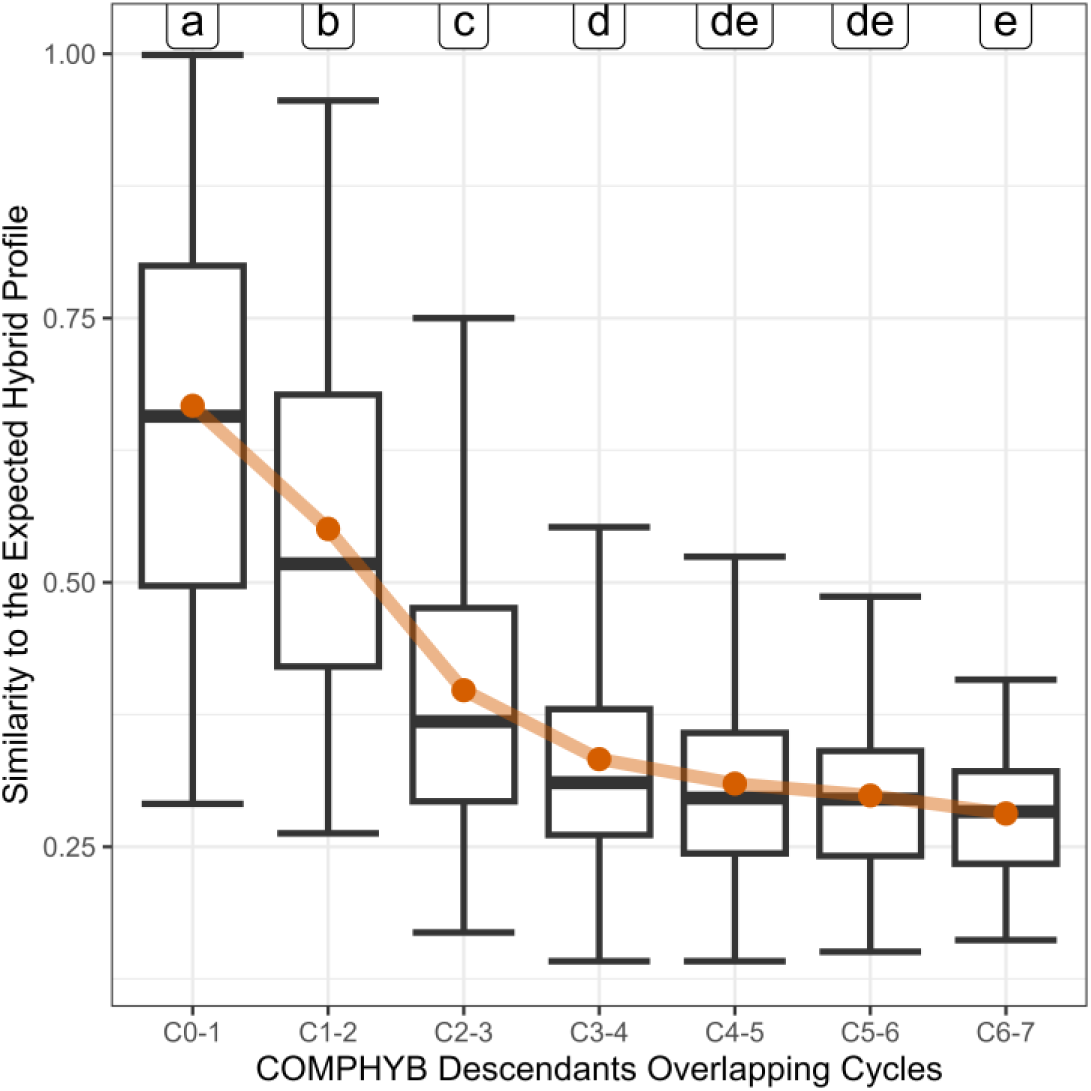
Evolution of the proportion of COMPHYB haplotypes monitored across its different progeny selection cycles. Boxplots of the haplotypic similarity between inbreds and the constructed hybrid profile. Orange dots represent the average similarity. Significance letters were obtained from Tukey HSD test.

### Expected and observed residual inbreeding in intergroup and intragroup crosses in dataset D2

We investigated the expected residual inbreeding of all possible parental crosses for the second dataset, representing a total of 106,491 pairwise potential single-cross hybrid combinations. This analysis was carried out to assess the differences between intragroup and intergroup crosses, and the possible difference between potential crosses and tested crosses. For all potential crosses, Pearson correlation between IBS and RIB was 0.99 (p-value < 2.2E-16), and application of a linear model explaining RIB with IBS (i.e., *RIB* = *IBSx* + *b*) indicated a beta effect of 1.962 and an intercept of -1.094. Consistently, RIB had a larger variance than IBS. Both RIB and IBS revealed significant difference between intergroup (mean[RIB] = 0.125; mean[IBS] = 0.620) and intragroup crosses (mean[RIB] = 0.328; mean[IBS] = 0.727) (p-value < 2.2E-16). However, a large overlap of the distributions was observed, notably in the low-range values (Supplementary Fig. 6). Segmentation of the intergroup and intragroup hybrids into assigned genetic group(s) revealed finer contrasts in residual inbreeding (Table 3 - Supplementary Fig. 7). Overall, RIB and IBS variances for intragroup crosses were consistently superior to intergroup crosses. For intragroup crosses, intragroup ADMX crosses were significantly superior in RIB with an average of 0.416. This value indicates that, on average, 42% of haplotypes are shared between 2 ADMX inbreds. For intergroup crosses, pairs with an ADMX inbred had significant superior intergroup parental similarity than the others (mean[SS-ADMX] = 0.181; mean[ADMX-IDT] = 0.211). Potential crosses between SS and IDT had the lowest average RIB similarity (0.086) and lowest RIB variance (0.031).

Further characterization of SS x IDT crossing pattern identified changes specific to the 97 hybrids found in the first dataset. The subset of 580 hybrids produced (D2HG) was compared to the set of all possible intergroup SS x IDT hybrid combinations (D2HP) and to the overall set of tested hybrids in 2024 (D2HM). According to both parental similarities’ methods, D2HP and D2HG did not have a significant difference in parental similarity (p-value[RIB] = 0.7224; p-value[IBS] = 0.4642). However, D2HM had significantly lower parental similarity than both D2HP and D2HG hybrids (Supplementary Fig. 8). Average genome-wide RIB was 12.36%, 12.52%, and 10.66%, for D2HP, D2HG and the 2024 D2HM hybrids, respectively. Genome-wide RIB mapping displayed few differences between D2HP, D2HG, and the D2HM, with similar hotspots for residual inbreeding (Fig. 5). The most striking contrast is the sharp significant loss of RIB on chromosome 10 for the D2HM (p-value = 2.5E-05), which went from 24.11% for global D2HG hybrids to 12.09% for 2024 ones.

**Fig. 5.**
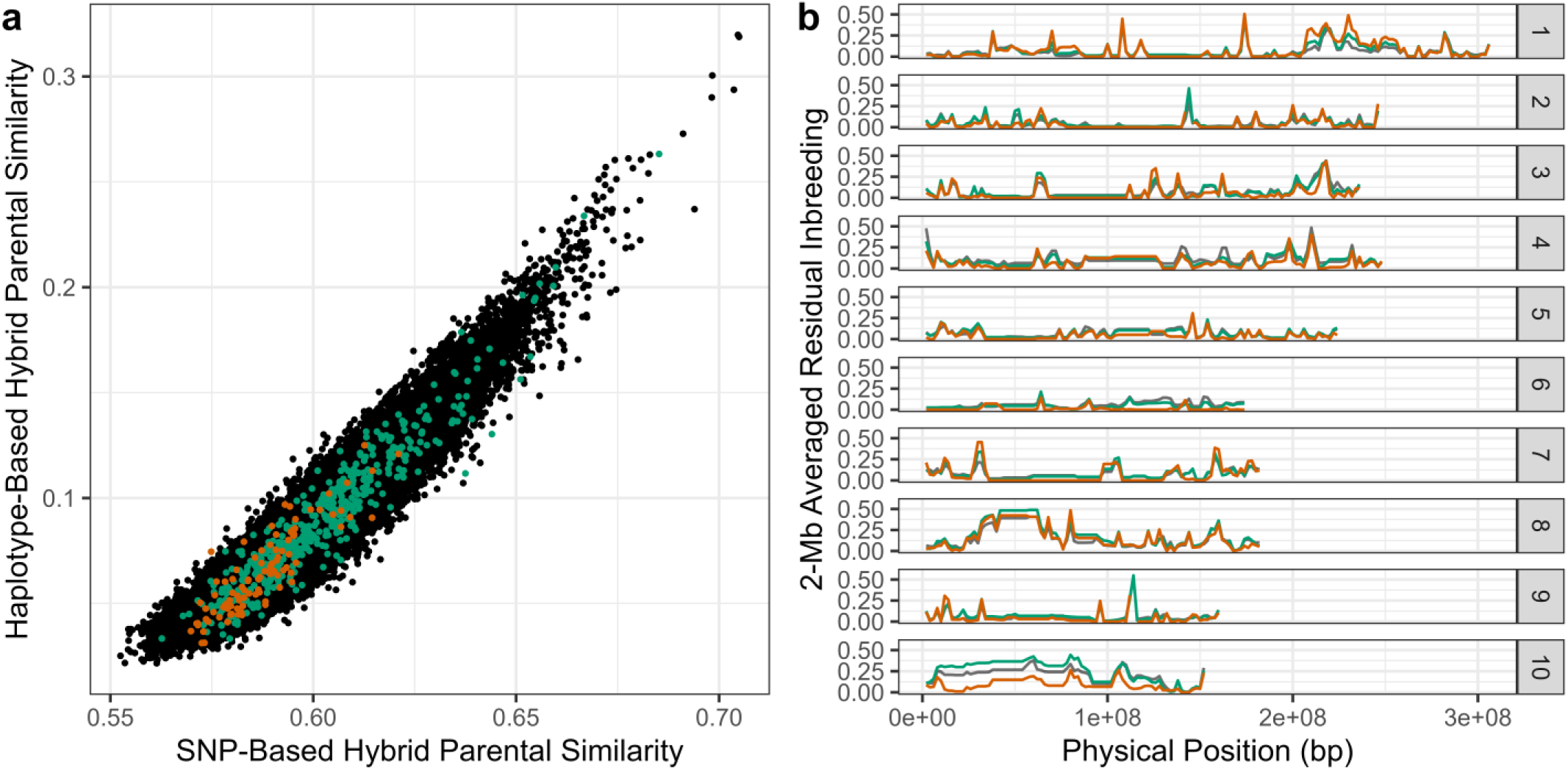
Overview of the expected hybrids’ residual inbreeding, obtained from the potential pairwise SS x IDT crosses from the available inbreds. a) Scatter plot between IBS and RIB. b) Evolution of RIB along the genome. Color scheme separates the potential SS x IDT hybrid crosses D2HP (black), from the global hybrid crosses D2HG (green) and the subset of modern 2024 hybrid crosses D2HM (orange).

### Transferred haplotypes origins and evolution across selection cycles in dataset D2

Assignation of the complete haplotypic dataset to specific origins, according to inbred year of availability, contributed to the characterization of residual inbreeding in the 580 tested SS x IDT intergroup hybrid crosses. Analysis of the origin-specific tested hybrids’ residual inbreeding highlighted two major regions: a Stiff Stalk-assigned fragment on chromosome 8 at 50 Mb transferred in Iodent material and an Iodent-assigned fragment close to chromosome-long on chromosome 10, transferred to Stiff Stalk inbreds (Fig. 6). Overall, observed residual inbreeding was heavily assigned to a Stiff Stalk origin. From the two major outlier fragments, we further categorized the origin and frequency evolution of their most common haplotypes.

**Fig. 6.**
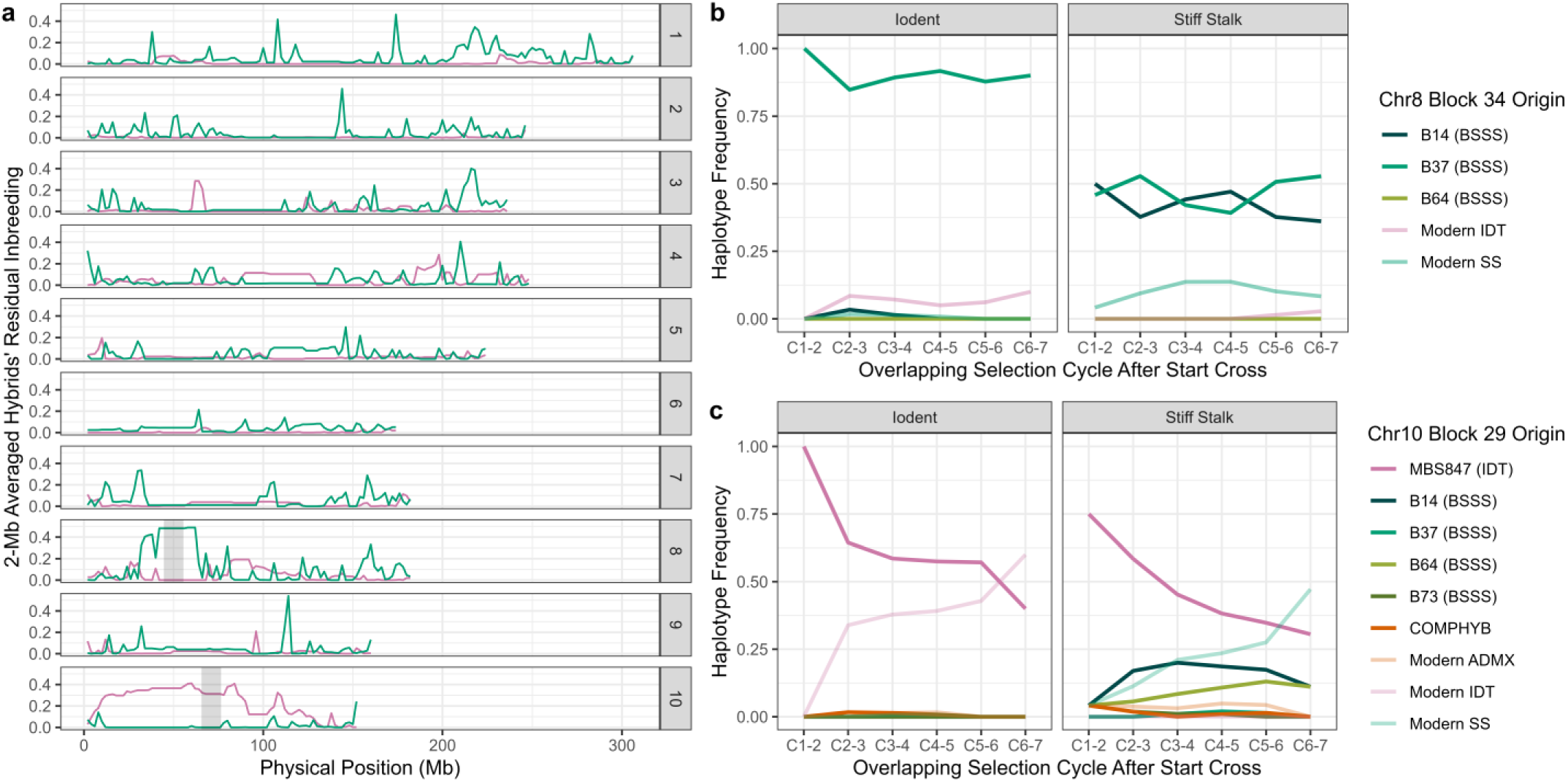
Decomposition of tested SS x IDT hybrids’ (D2HG) residual inbreeding and origin-specific characterization. a) Genome-wide residual inbreeding according to the haplotype genetic group origin (Green: SS and Pink: IDT). Grey zones on chromosomes 8 and highlight regions presented on graphs b and c. b) Evolution of haplotypic frequencies in Stiff Stalk and Iodent heterotic according to the overlapping selection cycle for the Chromosome 8 Haplotype Block 34 (CHR8B34). c) Evolution of haplotypic frequencies in Stiff Stalk and Iodent heterotic according to the overlapping selection cycle for the Chromosome 10 Haplotype Block 29 (CHR10B29). Color scheme for b) and c) correspond to the haplotype’s first assigned inbred. Haplotypes in b) and c) are ordered by global frequency: the top group reference haplotype (e.g., B14 in the CHR8B34) is H1, etc.

On chromosome 8, we focused on the haplotype block 34, located between 39.55 Mb and 55.93 Mb. This block has one of the highest RIBs of the chromosome, reaching 47.76%, while still being sufficiently long to limit spurious IBS (19 SNPs and 8 cM) . On this block, we detected 16 haplotypes, of which the two most frequent ones explain 89.07% of the dataset (B34H1 = 68.40% ; B34H2 = 20.67%). Group-specific frequencies exposed a clear contrast between these two foremost haplotypes. The haplotype B34H1 is found in 85.71% of ADMX inbreds and 89.5% of IDT inbreds, while the haplotype B34H2 is absent in both groups. In the SS group, the haplotypes B34H1 (43.69%) and B34H2 (44.66%) show similar frequencies. According to their year of access, the haplotype B34H1 is assigned to the BSSS reference B37 but is also carried by the Iodent group check MBS847, and all the COMPHYB selfings (SELF1 to SELF4), while the haplotype B34H2 was first found in B14 and consequently observed in B73, A632, the LG-B73 parent, and TC1. Monitoring of the haplotypes’ frequencies over the selection cycle confirmed the expected residual inbreeding in the hybrids, with stable frequencies for both B34H1, in the two groups, and B34H2, in the Stiff Stalks (Fig. 6). Proportion of homozygosity for B34H1 in 2024 tested hybrids reached 42.27%

On chromosome 10, we focused on the haplotype block 29. This block is 83 SNP markers long (i.e., 8.3 cM), has 59 recorded haplotypes, and is located between 65.36 Mb and 77.46 Mb. Haplotype frequency distribution is highly heterogeneous, with the most common haplotype (B29H1) found in 53.46% of inbreds, followed by the second most frequent (B29H2) at 7.36%. The distribution of B29H1 per inbreds’ genetic group was 82.14% for ADMX, 58.0% for IDT, and 43.69% for SS. Assignment of the earliest reference inbred indicated that this haplotype is unique to MBS847 and was propagated germplasm-wide through 4 of the 6 cycle 0 inbreds (TC1, TC2, SELF2, and SELF4). Evolution of haplotype frequencies highlighted a continuous decline in MBS847 haplotype (B29H1) in these materials, replaced by another IDT-specific haplotype in the Iodent group and another SS-specific haplotype in the Stiff Stalk group (Fig. 6). While leading to homozygous pairs in hybrids, this haplotype remained at high frequencies in both groups, even after 7 breeding cycles (40% in IDT and 31% in SS at C6-7). Conversely, the proportion of homozygosity for B29H1 in 2024 tested hybrids confirms this decline with only 7.99% of hybrids concerned.

## DISCUSSION

In the present study, we introduced a new haplotype-based approach to assess hybrid residual inbreeding using parental similarity. Subsequently, we characterized the main genetic origin that led to residual inbreeding in modern experimental elite hybrids from *Limagrain* mid-early/mid-late temperate maize breeding program.

### Relationship between parental similarity and hybrid performances

Knowledge of genetic relationships among inbreds is important to improve the organization of breeding programs, specifically in hybrid breeding schemes. Monitoring the genetic similarity among parents of intergroup hybrids can be a powerful tool to verify the divergence between heterotic groups and limit the risks of residual inbreeding (Bernardo 1993). The prediction of single-cross hybrid performances from molecular marker relationships to maximize performances and streamline intergroup crosses is thus a long-lasting goal for plant breeders (Bernardo 1992). Previous studies have demonstrated the correlation between inbreeding and heterosis for intragroup crosses, admixed or related materials (Burstin and Charcosset 1997; Larièpe et al. 2012; Larièpe et al. 2017, Roth et al., 2022). However, this correlation weakens with the analysis of intergroup hybrids, possibly linked with lower overall inbreeding values interval, low variance and different linkage disequilibrium profiles (Charcosset and Essioux 1994). For the SNP-based IBS method, our results are consistent with previous studies. In modern experimental hybrids, we found that increasing the similarity between parents by 1% reduced hybrids’ grain yield by -0.48 qt/ha when using SNP data. This regression coefficient is within the range of previously reported effects (Ramstein et al. 2020: -0.69 qt/ha per 1%; Roth et al. 2022 : between -1.53 and -0.21 qt/ha per 1%).

In this study, we proposed a parental relationship indicator based on haplotypes, which is significantly associated with maize grain yield and well adapted to trace back specific founder contributions. We compared this haplotype-based indicator (RIB) with a biallelic raw SNP approach (IBS). We found a more significant association between this haplotype-based indicator and yield, but the regression coefficient was lower (to -0.30 qt/ha for every 1%). The modification in effect can be explained by the different scales of the two indicators, where RIB varied between 0.02 and 0.18, while IBS varied between 0.55 and 0.64. Analyses using SNP biallelic data cannot differentiate similarities inherited by true ancestral relationships (i.e., presence of a pedigree relationship) from identity-by-state, generated from rare specific polymorphism, which do not give a better understanding of relationships (e.g., polymorphism specific to the separation between selected and wild types), mutations, or errors (e.g., genotyping, imputation). Utilization of multiallelic methods aims at reducing this proportion of unwanted IBS and considers more recent ancestry (Kim and Yoo 2016). While reported previously as non-significant for genomic prediction (Ramstein et al. 2020), incorporation of fixed residual inbreeding effect using this new method could positively improve prediction accuracy and will be presented in a future study. This new approach could notably affect prediction accuracy for hybrids whose both parents were never previously tested.

Application of these beta coefficients to extreme inbreeding cases (i.e., intragroup crosses and self-reproduction) suggests that pure-line inbreds, which have 100% residual inbreeding, should have grain yield averaging around 90-100 qt/ha. This theoretical yield appears unlikely high, and three hypotheses can be proposed to explain this discrepancy. First, this may suggest that additional effects, notably negative epistatic effects between homozygotes modify this linear trend to another (e.g., polynomial, exponential, logarithmic), further degrading inbred yield as similarity increases. Second, in this study, we used the commercial genotyping array of Limagrain Field Seeds. This genotyping method is used to differentiate and be highly polymorphic notably between heterotic groups in hybrid breeding. This can limit the observation of monomorphisms and rare alleles, leading to an ascertainment bias. Following Charcosset and Essioux (1994) recommendations, increasing the number of markers or using sequencing technology could also result in better correlation and significance, by having a better estimation of IBD and by being more appropriate to estimate inbreeding effect on phenotypic traits. It is also important to note that the use of a different haplotype construction method, with a different marker data preprocessing (e.g., LD pruning, MAF filtering) could have substantial consequences on the computation of the residual inbreeding, and may thus modify the value of the indicator and the value of the regression coefficient (Kim and Yoo 2016). Lastly, this projected inbred yield may be its true value. Although studies on inbred lines from commercial programs are rare, it has been found that the obtained genetic gain and increase in hybrid maize grain yield was associated with a parallel improvement of its parental inbreds, leading to a decrease in the percentage of heterosis (Troyer 2006).

The comparison between intragroup and intergroup parental similarity distributions exposed significantly lower values for intergroup crosses, which highlights that the current management practice relying on divergent crossing populations is efficient. However, a partial overlap between intra-and intergroup parental similarities can still be noticed. The intragroup Stiff Stalk distribution had the largest overlapping distribution tail, with 4% of intragroup SS inbred pairs having lower residual inbreeding than the 3^rd^ quantile distribution of intergroup pairs’ residual inbreeding (Supplementary Fig. 7). High dissimilarity between intragroup inbreds demonstrates the potential for high-performing hybrids obtained from originally similar genetic backgrounds (Bernardo 2001). Application of this method could be used to decipher sub-heterotic groups in the germplasm. Further decomposition of the population structure may be useful to either understand more precise crossing patterns or select more representative tester inbreds for intergroup evaluation.

### Maize flowering and germplasm-wide conservation of haplotypes

Our analysis identified two haplotypes within the progeny of the competitor hybrid start crosses that persisted at high frequencies in the two opposite heterotic groups. While leading to significant residual inbreeding in hybrids, these fragments were preserved across selection cycles. In the prospect of maximizing heterosis and heterotic group divergence, long-lasting retention of undesirable haplotypes may underline issues in the breeding pipeline. However, both observed haplotype block regions are located on chromosomes 8 and 10, consistent with previous studies on temperate maize flowering. Flowering time is a critical issue for plant breeding, as it limits the cultivation area and the logistics around the two crossing pools (i.e., flowering interval between female and male inbreds) and impacts hybrids’ yield and harvest quality (Ducrocq et al. 2009). While primarily quantitative, flowering-derived traits (e.g., day to anthesis, day to silking, anthesis-silking interval, etc.) are linked to major quantitative trait loci (QTL) with clearly defined effects. Numerous studies have localized high-impact regions on chromosome 8 and chromosome 10, related to the loci *vgt1*, *ZmCCT*, *ZFL*, and other QTL hotspots (Chardon et al. 2004; Buckler et al. 2009; Ducrocq et al. 2009; Gouesnard et al. 2017; Rio et al. 2020).

It is to be noted that while we focused on only 2 haplotype blocks (Fig. 6), origin-specific residual inbreeding regions were significantly larger, particularly for chromosome 10, where it encompasses most of the physical length of the chromosome. This high retention of Iodent germplasm inside the Stiff Stalk heterotic group may be related to the reported significantly higher LD and longer haplotype length for this chromosome (van Heerwaarden et al. 2012; Schaefer and Bernardo 2013; Romay et al. 2013; Wu et al. 2016). Higher homozygosity in flowering-related regions coincides with demonstrated mainly additive or partially dominant effects, which may prompt the necessity of stacking recessive alleles in hybrid profiles (Ducrocq et al. 2008; Ducrocq et al. 2009). This compulsory homozygosity was also revealed to be in most cases origin-specific (Romero-Severson et al. 2001; Schaefer and Bernardo 2013; Rio et al. 2020). Lauer et al. (2012) notably showed consistent earlier flowering dates with the introgression of Minnesota 13-type and Iodent inbreds in single-cross hybrids. Earlier maturity in hybrids is notably one of the reasons linked with the current over-representation of Iodent material in breeding schemes (Smith 2007). Yet, while we cannot fully exclude that observed frequency evolution is solely due to genetic drift, we hypothesize that similar haplotype retention in both heterotic groups on chromosome 8 and 10 is linked with earlier precocity needed to adapt and better perform in European colder climatic conditions.

### Impact of the introgression of novel material in hybrid recurrent reciprocal breeding schemes

Modern maize hybrid breeding relies on the systematic crosses of inbred parents from different divergent heterotic groups (Melchinger et al. 1991; Mikel 2008). This stringent management practice helps to maximize heterosis, limits residual inbreeding, and consequently contributes to the significant superior performances of hybrid breeding (Penny and Eberhart 1971; Crow 1998). The global dissemination of RRS-type breeding approaches in public and private breeding schemes and the reported small maize founder population size for temperate germplasm supported the reproductive isolation between heterotic groups and the heavy recycling of top-performing inbred lines for both intragroup development, as start-cross material, and as testers to evaluate the opposite group, for intergroup crosses (Smith et al. 2004; Reif et al. 2005; Smith et al. 2022). The restriction of inbred development to intragroup crosses and the global dependence on common elite material cause a reduction in genetic diversity, lowering the effective population size, and lead to the fixation of different genomic regions in each group (Allier et al. 2019; Smith et al. 2022), notably in *Limagrain* dent maize germplasm (Kadoumi et al. 2025). Genetic diversity management is subsequently an essential aspect of plant breeding to achieve constant genetic gain and mitigate the risks associated with the repeated selection of elite material (Technow et al. 2021). However, germplasm diversification, the introduction of novel material, and its management are challenging tasks, as they may disrupt the divergent population structure. Here, we highlight the risks related to the incorporation of intergroup admixed material in a highly differentiated breeding scheme organized around already defined heterotic groups. We also highlight the potential utility and justification of superior inbreeding in specific genomic fragments. While the haplotype responsible for residual inbreeding in hybrids could have been seen as additional germplasm diversity, its purification and replacement by a heterotic group-specific modern haplotype did not seem to have a negative impact on performance. This decline in available haplotypes and genetic diversity thus appears to be desirable.

The utilization of commercial competitor hybrids as parents for inbred development is a distinctive approach to introgress state-of-the-art modern elite genetic backgrounds. Compared to other sources (e.g., exPVPs, landraces, etc.), which can be underperforming due to the time elapsed since their creation or lack of fitness, this genetic resource enables to bring very recent elite diversity expected to bring new top performing favorable alleles, notably for grain yield (Raposo et al. 2004; Reis et al. 2009; Rodrigo et al. 2012; Reis et al. 2014; Beckett et al. 2017; Almeida et al. 2024). This method is distinctively possible in UPOV (i.e. the international Union for the Protection of New Varieties of Plants) member states thanks to current regulations on breeding schemes, intellectual property, and the breeder exemption to use competitor hybrid material for breeding. It is notably utilized in countries without a common and fixed heterotic population structure, and where the American dent x dent pattern cannot be applied, mostly due to precocity/tardiness issues, like China, Thailand, or Brazil (Ferreira et al. 2010; Oliboni et al. 2013; Senhorinho et al. 2015; Leng et al. 2019; de Faria et al. 2022; Smith et al. 2022; Luz et al. 2024). However, as seen in this article and independent of the selected breeding cross (i.e., selfing or topcross), this method can lead to the transfer of haplotypic regions between crossing pools. The creation of inbreds with a mixture of germplasm with different origins also presents logistical issue to allocate a highly complex genetic background with a good combining tester. This increased admixture between heterotic groups is linked to a decrease in divergence, an increased residual inbreeding of the intergroup hybrids which has a negative effect on hybrid performances. Our results show that the consequences of such an introduction in the 80s were still found in the breeding scheme after several cycles of selection. Introgression of exPVP material is a parallel consideration to this issue, which, due to intellectual property laws, will have a 20-year-long lag (or 25 years for trees and vines), and can impose a yield or quality penalty (Mikel 2011; Kurtz et al. 2016). However, this source of germplasm is better characterized, with substantial literature (Mikel and Dudley 2006; Mikel 2006; Mikel 2011; Kurtz et al. 2016), and lowers risks of bringing unwanted admixture into the selected group (Nelson et al. 2008; Guo et al. 2021). These differences in the modality of access to elite genetic variability are an ongoing topic in plant breeding, and notably maize hybrid breeding (Hao et al. 2020; Allier et al. 2020; Sanchez et al. 2023).

Multiple articles have proposed the introduction of admixed intergroup lines to manage effectively long-term genetic diversity (Bernardo 2001; Roth et al. 2022; Zhang et al. 2024; Bernardo 2025). However, while possible, the disruption of the long-established heterotic groups led to additional logistical issues, particularly finding suitable tester lines to evaluate the new inbreds in hybrid crosses (Bernardo 2001). This article highlights the consequences of modifying the current heterotic population structure management and supports the introgression of supplemental genetic diversity through other genetic resources, in the context of direct use in the inbred development pipeline. The careful introduction of useful tropical and landrace germplasm and the use of exPVPs material should thus remain privileged methods for diversity management in hybrid divergent breeding schemes (Smith et al. 2022). Nevertheless, as reported by Allier et al. (2020) and Roth et al. (2022), admixed lines could still be regarded as potential pre-breeding material.

## CONCLUSION

We presented a novel haplotype-based approach for assessing parental genetic similarity and its implications for evaluating residual inbreeding in temperate maize hybrids. Compared to traditional biallelic identity-by-state metrics, this multi-allelic residual inbreeding estimator demonstrated a significantly stronger association with hybrid grain yield in dent maize, providing a more precise tool for hybrid evaluation and ranking. The characterization of high residual inbreeding in hybrids supported the identification of shared genetic fragments across both groups of the heterotic pattern. This set of common haplotypes is derived to a large extent from the introduction of a commercial competitor F1 hybrid. This research thus represents the first published analysis of commercial hybrid use in private inbred breeding programs and its consequences on haplotype transfers between heterotic groups. The identification of population-wide persistent transfers, albeit in a downward frequency trend, underlines the potential risk of inbred development with intergroup crosses. Yet, observation of persisting haplotype similarity over multiple selection cycles, suggests a non-monitored positive selection pressure likely associated with early flowering. Careful management and monitoring are consequently necessary to benefit from such effects while mitigating the disruption of heterotic group divergence. This work highlights the importance of diversity management and in-depth germplasm knowledge in sustaining the heterotic population structure. By addressing the consequences of residual inbreeding, we support novel breeding strategies aimed at maintaining heterotic integrity and enhancing hybrid yield potential.

## AUTHOR CONTRIBUTION STATEMENT

NH, FH, MB, AM, AC and LM initiated and supervised the project. RK designed and ran the analyses, interpreted the results, and prepared the manuscript. NH, FH, MB, AM, AC, LM designed the study and assisted in the interpretation of the results. All authors contributed, revised and approved the manuscript.

## DATA AVAILABILITY

The data analyzed in this study are part of an active commercial maize breeding program of *Limagrain Field Seeds* and is composed of the latest experimental hybrids for its mid-early to mid-late maturity. As such, the phenotypic data, genotypic material and pedigree information are confidential and protected as intellectual property or as trade secrets. Consequently, datasets are not publicly available.

## COMPLIANCE

Conflict of interest

Authors AC and LM declare they have no conflict of interests. Authors RK, NH, FH, MB, AM are employed by *Limagrain Field Seeds*. This research was funded by *Limagrain Field Seeds* and the *Agence Nationale de la Recherche et de la Technologie (ANRT*) *[CIFRE Grant N°2023/1777]* for RK.

## ACKNOWLEDGEMENT

This research was funded by *Limagrain Europe* and the *Agence Nationale de la Recherche et de la Technologie* (ANRT) [CIFRE Grant N°2023/1777] for RK. The authors would like to thank *Limagrain Europe* for providing the molecular, pedigree and phenotypic data. The author RK wishes to thank everyone involved in the proofreading of this manuscript and the discussions around this project. We thank the members of the INRAE-CIRAD “R2D2” network for helpful discussions on genomic selection in breeding programs.

