## Supplementary Material for "Identification of heterotic group-specific haplotypes and impact of residual inbreeding on grain yield of maize elite hybrids"

Short Title: Inbreeding effect on maize hybrid yield

#### AUTHORS

Romain Kadoumi (https://orcid.org/0009-0007-0560-4945)[1,2], Nicolas Heslot[1], Fabienne Henriot[1], Alain Murigneux[1], Mathilde Berton[1], Laurence Moreau (https://orcid.org/0000-0002-7195-1327)[2], Alain Charcosset (https://orcid.org/0000-0001-6125-503X)[2]

[1] Limagrain Field Seeds, 28 Route d'Ennezat, 63720 Chappes, France

[2] Université Paris-Saclay, INRAE, CNRS, AgroParisTech, Génétique Quantitative et Evolution (GQE) - Le Moulon, 91190, Gif-Sur-Yvette, France

**Corresponding author**

#### KEY MESSAGE:

Haplotype-based hybrid's parental similarity better predicts grain yield performance than IBS. Utilization of commercial hybrids as inbred start material resulted in higher hybrid residual inbreeding even after several selection cycles.

**KEY WORDS:**

Maize, Inbreeding, Haplotype, Genomic model, Heterotic group

#### ACKNOWLEDGEMENT

This research was funded by *Limagrain Europe* and the *Agence Nationale de la Recherche et de la Technologie* (ANRT) [CIFRE Grant N°2023/1777] for RK. The authors would like to thank *Limagrain Europe* for providing the molecular, pedigree and phenotypic data. The author RK wishes to thank everyone involved in the proofreading of this manuscript and the discussions around this project. We thank the members of the INRAE-CIRAD "R2D2" network for helpful discussions on genomic selection in breeding programs.

**This file contains:**

Supplementary Table Legends
Supplementary Table
Supplementary Figure Legends
Supplementary Figures

#### SUPPLEMENTARY TABLE LEGENDS

### SUPPLEMENTARY TABLES

***Supplementary Table 1** Distribution of inbreds per genetic group and per selection cycle*

| | REFERENCE | C0 | C1 | C2 | C3 | C4 | C5 | C6 | C7 |
|---|---|---|---|---|---|---|---|---|---|
| ADMIXED | 0 | 5 | 13 | 16 | 15 | 5 | 2 | 0 | 0 |
| IODENT | 1 | 0 | 0 | 4 | 55 | 85 | 35 | 14 | 6 |
| STIFF STALK | 6 | 1 | 8 | 16 | 37 | 58 | 44 | 25 | 11 |

***Supplementary Table 2** Pairwise comparisons of the sixteen models according to the difference in AIC and BIC, and the Chi-squared Likelihood Test p-value*

| | | | Chi-squared Likelihood Ratio Test P-value | | | | | | | | | | | | | | | |
|---|---|---|---|---|---|---|---|---|---|---|---|---|---|---|---|---|---|---|
| | | | Null Model | | | | Model with IBS | | | | Model with RIB | | | | Model with IBS & RIB | | | |
| | | | A | AD | A(AA) | AD(AA) | A | AD | A(AA) | AD(AA) | A | AD | A(AA) | AD(AA) | A | AD | A(AA) | AD(AA) |
| Delta Difference in AIC | Null Model | A | - | 6.4E-13 | 2.8E-15 | 2.4E-15 | | | | | | | | | | | | |
| | | AD | 49.7 | - | - | 7.8E-05 | | | | | | | | | | | | |
| | | A(AA) | 60.4 | 10.7 | - | 2.6E-02 | | | | | | | | | | | | |
| | | AD(AA) | 63.3 | 13.6 | 2.9 | - | | | | | | | | | | | | |
| | Model with IBS | A | 13.8 | -35.8 | -46.6 | -49.5 | - | 2.1E-12 | 9.3E-15 | 8.2E-15 | | | | | | | | |
| | | AD | 61.2 | 11.5 | 0.8 | -2.1 | 47.4 | - | - | 8.4E-05 | | | | | | | | |
| | | A(AA) | 71.9 | 22.2 | 11.5 | 8.6 | 58.0 | 10.6 | - | 2.8E-02 | | | | | | | | |
| | | AD(AA) | 74.7 | 25.0 | 14.3 | 11.4 | 60.9 | 13.5 | 2.8 | - | | | | | | | | |
| | Model with RIB | A | 15.7 | -34.0 | -44.7 | -47.6 | 1.9 | -45.5 | -56.2 | -59.0 | - | 4.2E-12 | 2.6E-14 | 2.2E-14 | | | | |
| | | AD | 61.7 | 12.0 | 1.4 | -1.6 | 47.9 | 0.5 | -10.1 | -13.0 | 46.0 | - | - | 1.2E-04 | | | | |
| | | A(AA) | 71.7 | 22.0 | 11.3 | 8.4 | 57.9 | 10.5 | -0.2 | -3.0 | 56.0 | 10.0 | - | 2.7E-02 | | | | |
| | | AD(AA) | 74.6 | 24.9 | 14.2 | 11.3 | 60.8 | 13.4 | 2.7 | -0.1 | 58.9 | 12.9 | 2.9 | - | | | | |
| | Model with IBS & RIB | A | 20.5 | -29.2 | -39.9 | -42.8 | 6.6 | -40.7 | -51.4 | -54.2 | 4.8 | -41.3 | -51.2 | -54.1 | - | 3.2E-12 | 2.0E-14 | 1.6E-14 |
| | | AD | 67.1 | 17.3 | 6.7 | 3.7 | 53.2 | 5.8 | -4.8 | -7.6 | 51.3 | 5.3 | -4.7 | -7.5 | 46.6 | - | - | 1.1E-04 |
| | | A(AA) | 77.0 | 27.3 | 16.6 | 13.7 | 63.2 | 15.8 | 5.1 | 2.3 | 61.3 | 15.3 | 5.3 | 2.4 | 56.5 | 9.9 | - | 2.6E-02 |
| | | AD(AA) | 78.0 | 30.3 | 19.6 | 16.7 | 66.2 | 18.8 | 8.1 | 5.3 | 64.3 | 18.2 | 8.3 | 5.4 | 59.5 | 12.9 | 3.0 | - |

*Supplementary Table 3* Evolution of origin-specific haplotypic similarities inside the population

| Cycle | | Avg. | 0 – 1 | 1 – 2 | 2 – 3 | 3 – 4 | 4 – 5 | 5 – 6 | 6 – 7 |
|---|---|---|---|---|---|---|---|---|---|
| **BSSS** | **Global** | 38.9 ± 24.7 | 49.8 ± 17.9 | 48.2 ± 20.2 | 39.0 ± 23.9 | 35.3 ± 25.3 | 36.6 ± 25.8 | 40.9 ± 24.8 | 43.0 ± 23.9 |
| | **SS** | 64.2 ± 9.41 | 70.4 ± 8.99 | 68.8 ± 7.07 | 67.1 ± 6.98 | 66.0 ± 8.84 | 63.8 ± 9.92 | 61.0 ± 9.37 | 59.7 ± 9.19 |
| | **IDT** | 14.0 ± 3.46 | - | 19.4 ± 2.58 | 15.5 ± 3.36 | 14.3 ± 3.50 | 13.4 ± 3.33 | 12.8 ± 2.81 | 12.8 ± 3.25 |
| | **ADMX** | 36.9 ± 9.95 | 39.4 ± 10.7 | 35.1 ± 10.8 | 35.6 ± 10.1 | 37.2 ± 8.49 | 36.5 ± 7.04 | 37.1 ± 8.68 | - |
| **MBS847** | **Global** | 25.1 ± 15.8 | 25.0 ± 11.4 | 25.7 ± 14.4 | 27.1 ± 15.1 | 27.1 ± 15.3 | 25.3 ± 16.3 | 21.6 ± 16.9 | 19.3 ± 16.4 |
| | **SS** | 9.07 ± 3.66 | 14.1 ± 2.41 | 12.0 ± 3.33 | 10.0 ± 3.38 | 9.36 ± 3.55 | 8.61 ± 3.68 | 7.82 ± 3.46 | 7.71 ± 2.90 |
| | **IDT** | 39.2 ± 7.77 | - | 43.9 ± 3.97 | 39.2 ± 8.06 | 38.6 ± 7.90 | 39.0 ± 7.78 | 40.4 ± 7.28 | 40.2 ± 7.13 |
| | **ADMX** | 32.5 ± 9.43 | 30.4 ± 10.0 | 34.6 ± 10.5 | 33.1 ± 9.83 | 30.8 ± 8.23 | 35.0 ± 4.73 | 34.7 ± 2.12 | - |
| **COMPHYB** | **Global** | 35.5 ± 15.1 | 66.7 ± 22.0 | 55.1 ± 17.4 | 39.8 ± 14.6 | 33.3 ± 10.8 | 31.0 ± 9.66 | 29.8 ± 8.41 | 28.1 ± 6.22 |
| | **SS** | 31.5 ± 9.19 | 42.9 ± 8.89 | 41.0 ± 8.87 | 34.6 ± 9.82 | 32.3 ± 8.83 | 30.3 ± 8.62 | 27.8 ± 7.14 | 27.5 ± 5.95 |
| | **IDT** | 30.7 ± 7.01 | - | 46.7 ± 9.46 | 33.3 ± 7.45 | 30.2 ± 6.65 | 29.6 ± 6.55 | 31.3 ± 6.32 | 29.3 ± 6.67 |
| | **ADMX** | 67.1 ± 15.2 | 78.6 ± 15.9 | 67.9 ± 13.3 | 61.1 ± 11.5 | 59.0 ± 9.59 | 63.6 ± 12.2 | 63.8 ± 18.4 | - |
| **UNKNOWN** | **Global** | 13.0 ± 3.35 | 7.42 ± 4.16 | 9.10 ± 3.28 | 11.9 ± 3.31 | 13.3 ± 2.80 | 14.0 ± 2.61 | 14.5 ± 2.20 | 14.8 ± 1.57 |
| | **SS** | 13.9 ± 2.52 | 11.5 ± 1.64 | 11.7 ± 1.83 | 13.0 ± 2.67 | 13.6 ± 2.69 | 14.2 ± 2.49 | 14.9 ± 1.93 | 14.9 ± 1.63 |
| | **IDT** | 13.8 ± 2.27 | - | 9.40 ± 2.23 | 13.0 ± 2.50 | 13.8 ± 2.30 | 14.1 ± 2.16 | 14.3 ± 1.95 | 14.5 ± 1.45 |
| | **ADMX** | 7.02 ± 3.06 | 5.40 ± 3.49 | 6.89 ± 2.69 | 7.98 ± 2.66 | 8.52 ± 2.11 | 6.93 ± 1.75 | 6.70 ± 2.81 | - |

#### SUPPLEMENTARY FIGURES

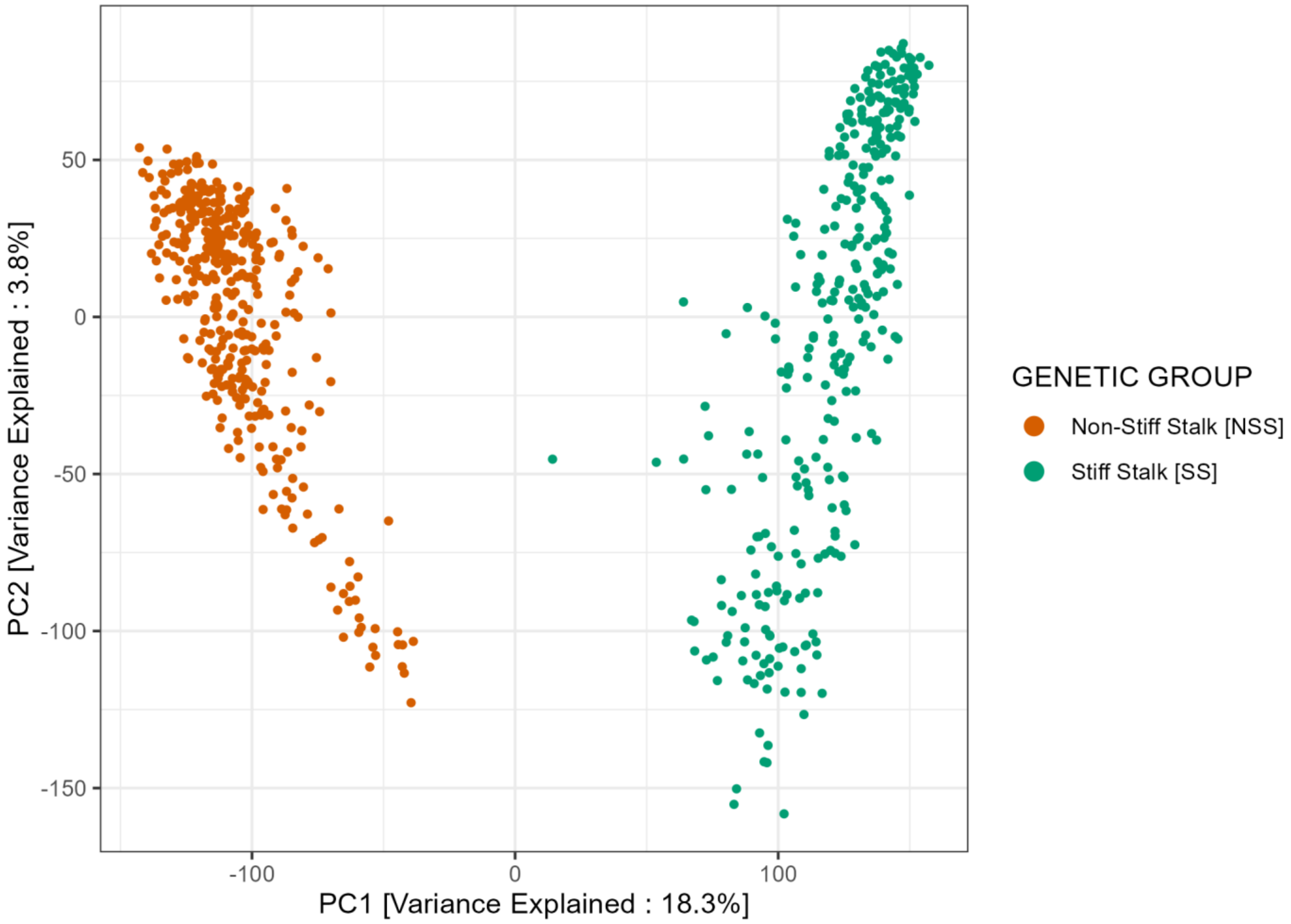

***Supplementary Fig. 1*** *Dataset 1 inbreds' population structure visualized by SNP-based PCA scatterplots (PC1 and PC2). Color scheme represents the inbred genetic group: Stiff Stalk (green) and Non-Stiff Stalk (orange)*

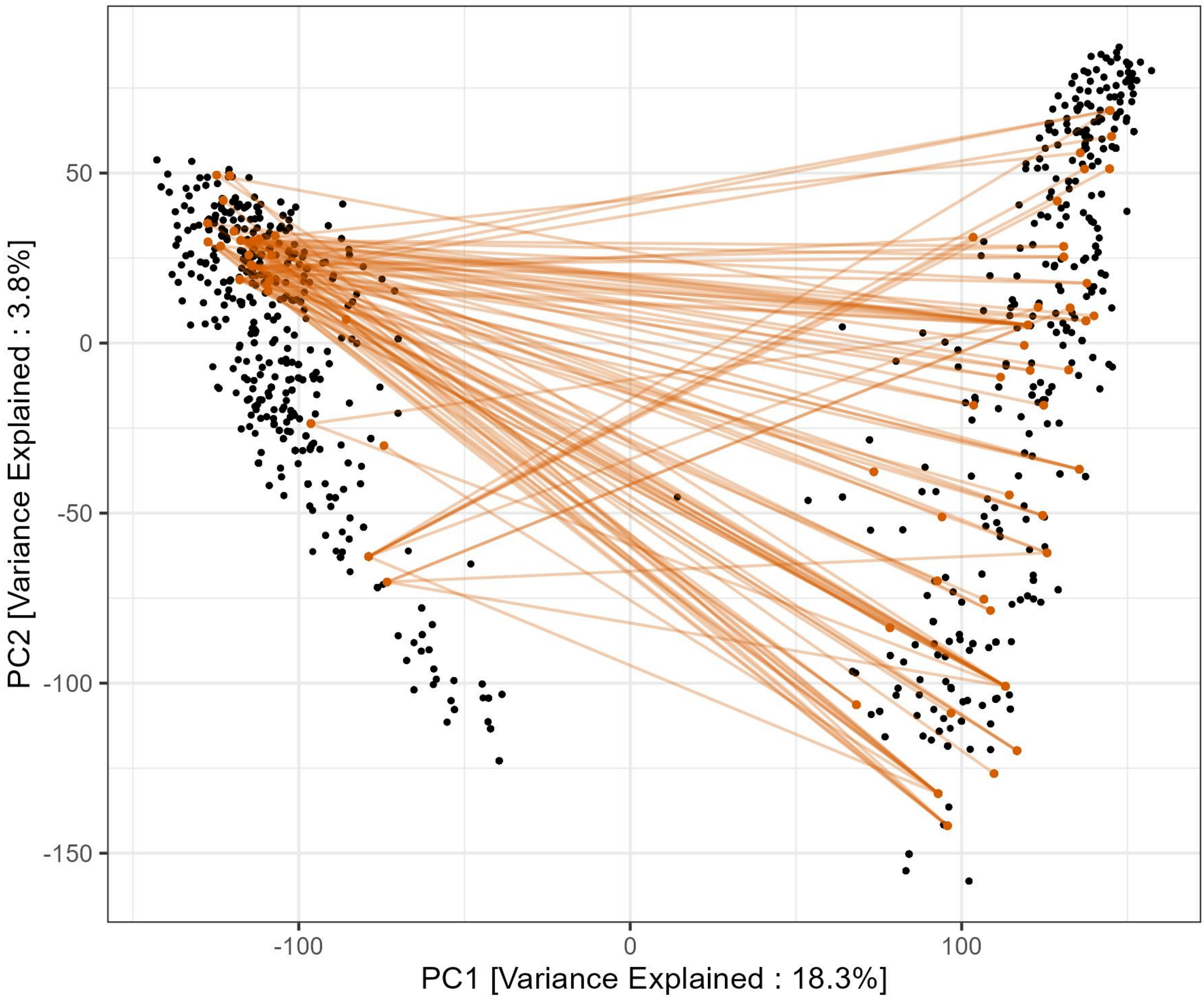

***Supplementary Fig. 2*** *Dataset 1 inbreds' population structure visualized by SNP-based PCA scatterplots (PC1 and PC2). Black dots indicate inbreds used in 2024 experimental hybrids. Orange dots indicate the inbred subset, originating from a common ancestor (COMPHYB). Orange lines link the parents of the 97 hybrids crosses.*

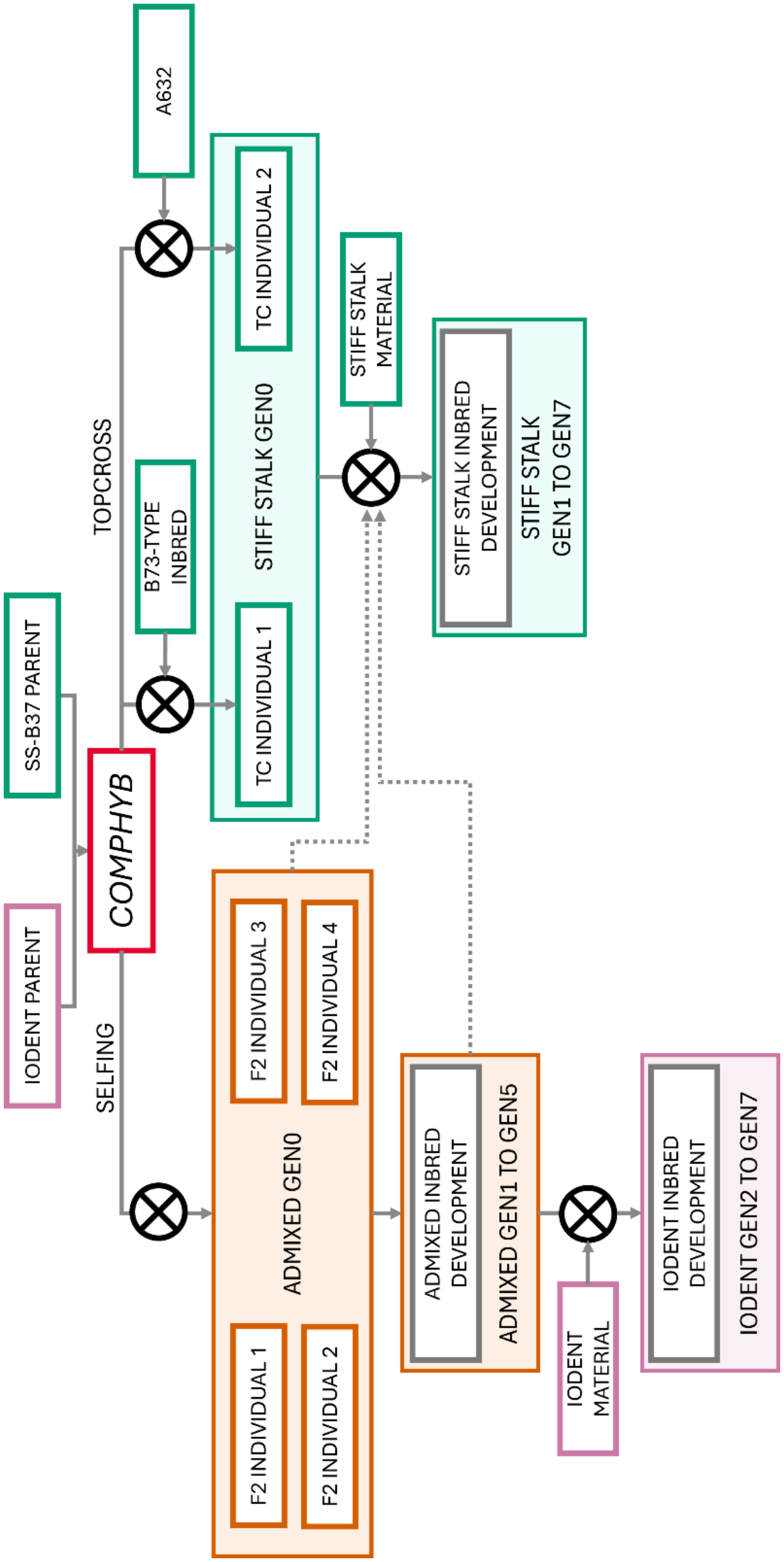

***Supplementary Fig. 3*** *Schematic representation of the pedigree of the inbred lines derived from the competitor commercial hybrid*

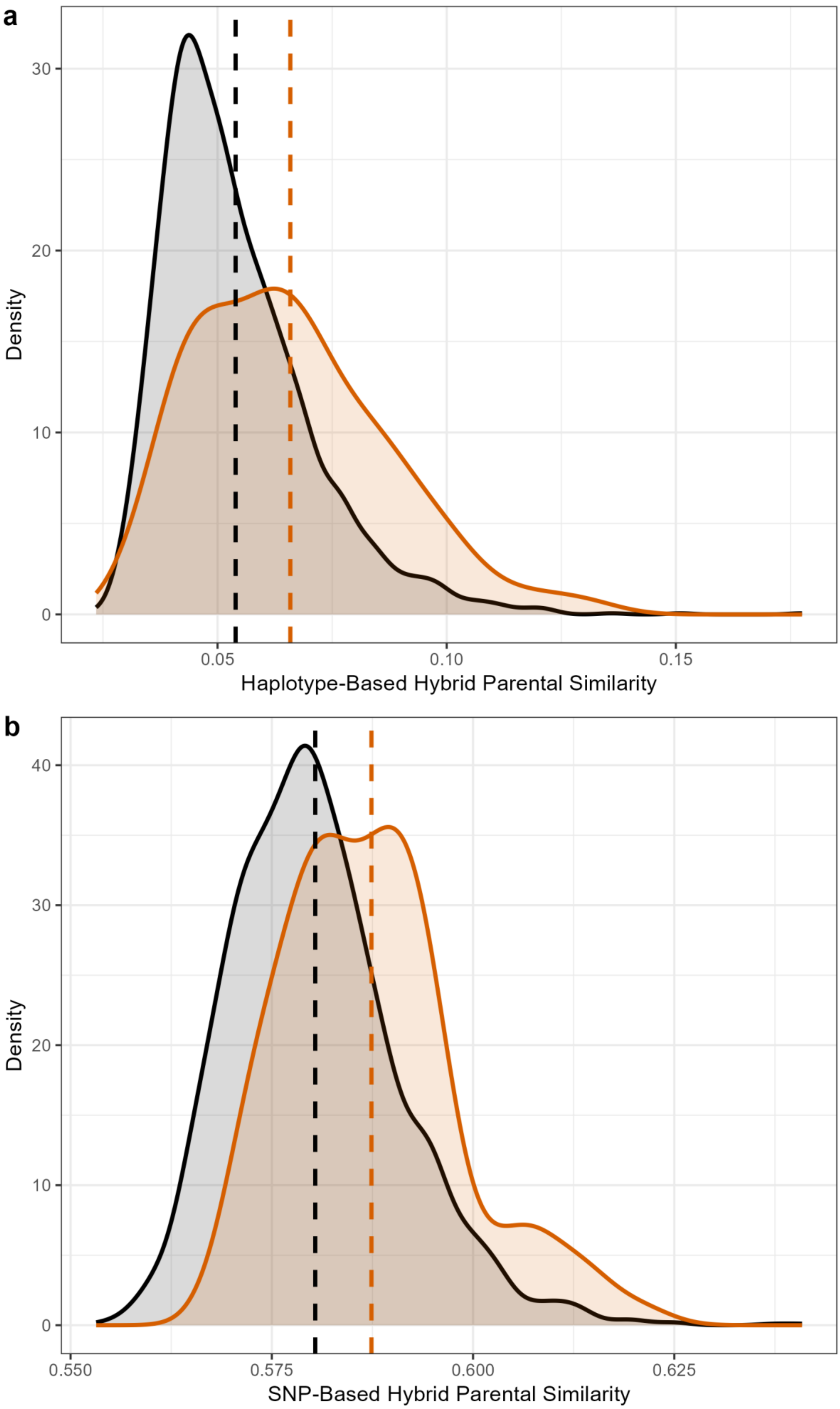

***Supplementary Fig. 4*** *Density diagrams of 2024 experimental hybrids' residual inbreeding measured by the haplotype-based method (RIB) (top) and SNP-based (IBS) (bottom). Vertical line represents the average of the specific category. Color scheme separates the global 2024 experimental hybrids dataset (black) to the hybrids with both parents originating from the competitor commercial hybrid inbred development (orange)*

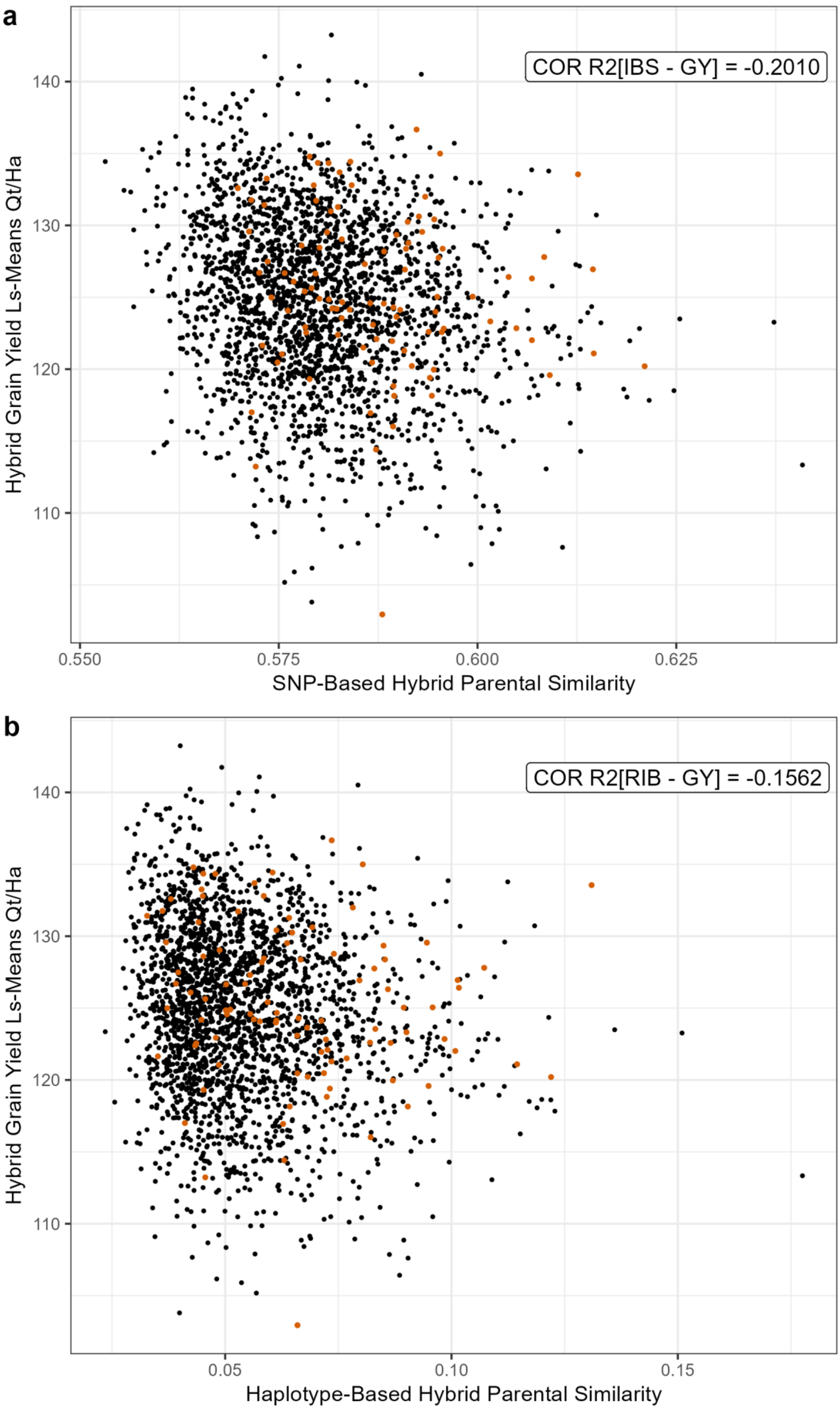

***Supplementary Fig. 5*** *Comparison between Hybrids' parental similarity indicators and Grain Yield lsmeans. Color scheme separates the global 2024 experimental hybrids dataset (black) to the hybrids with both parents originating from the competitor commercial hybrid inbred development (orange)*

85

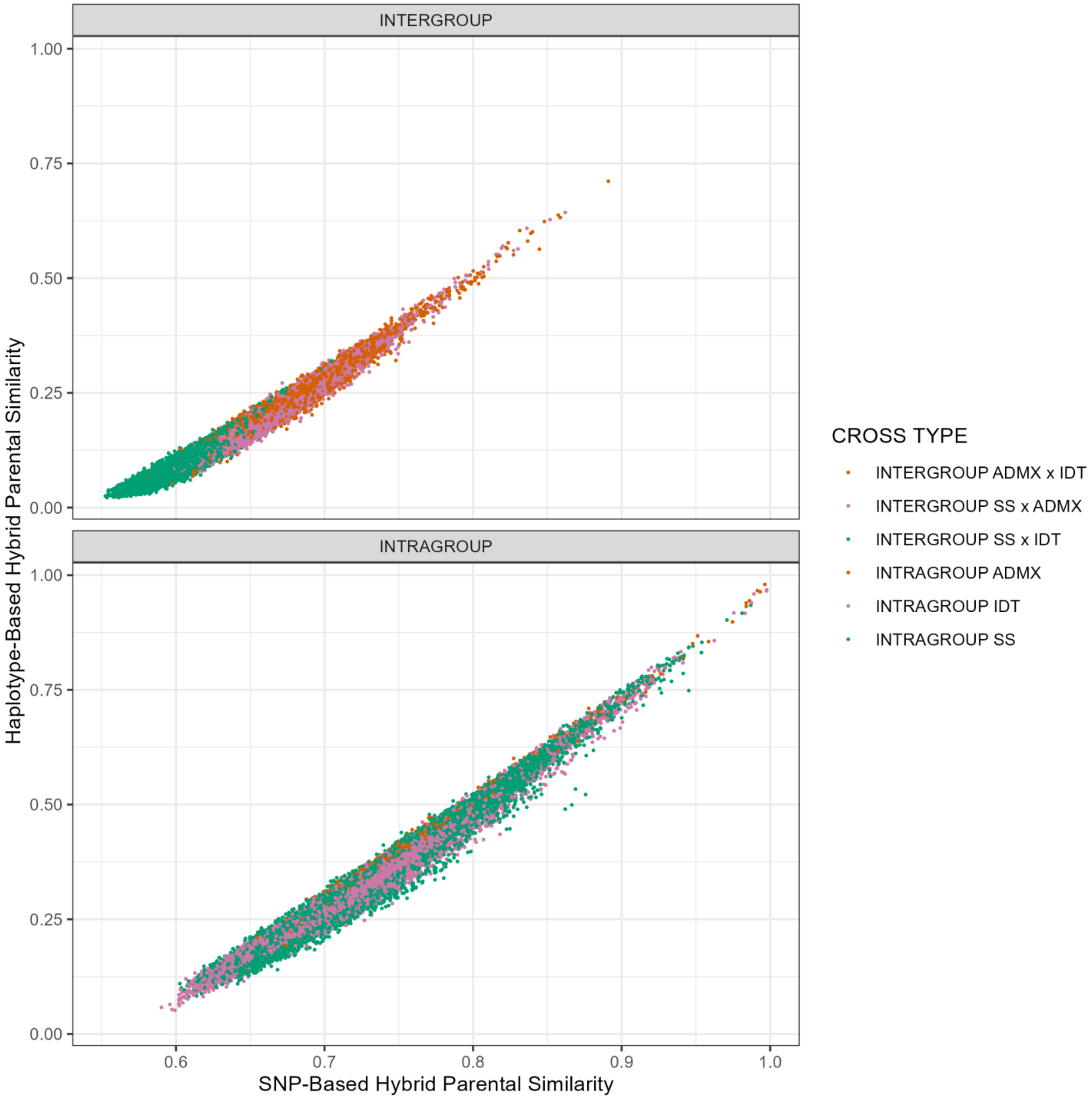

***Supplementary Fig. 6*** *Scatter plots of the SNP-based parental similarity indicator (IBS) compared to the haplotype-based one (RIB) for the complete possible pairwise combinations of all 462 inbreds linked to the competitor commercial hybrids inbred start cross. Scatter plots are separated according to the cross type (intragroup / intergroup)*

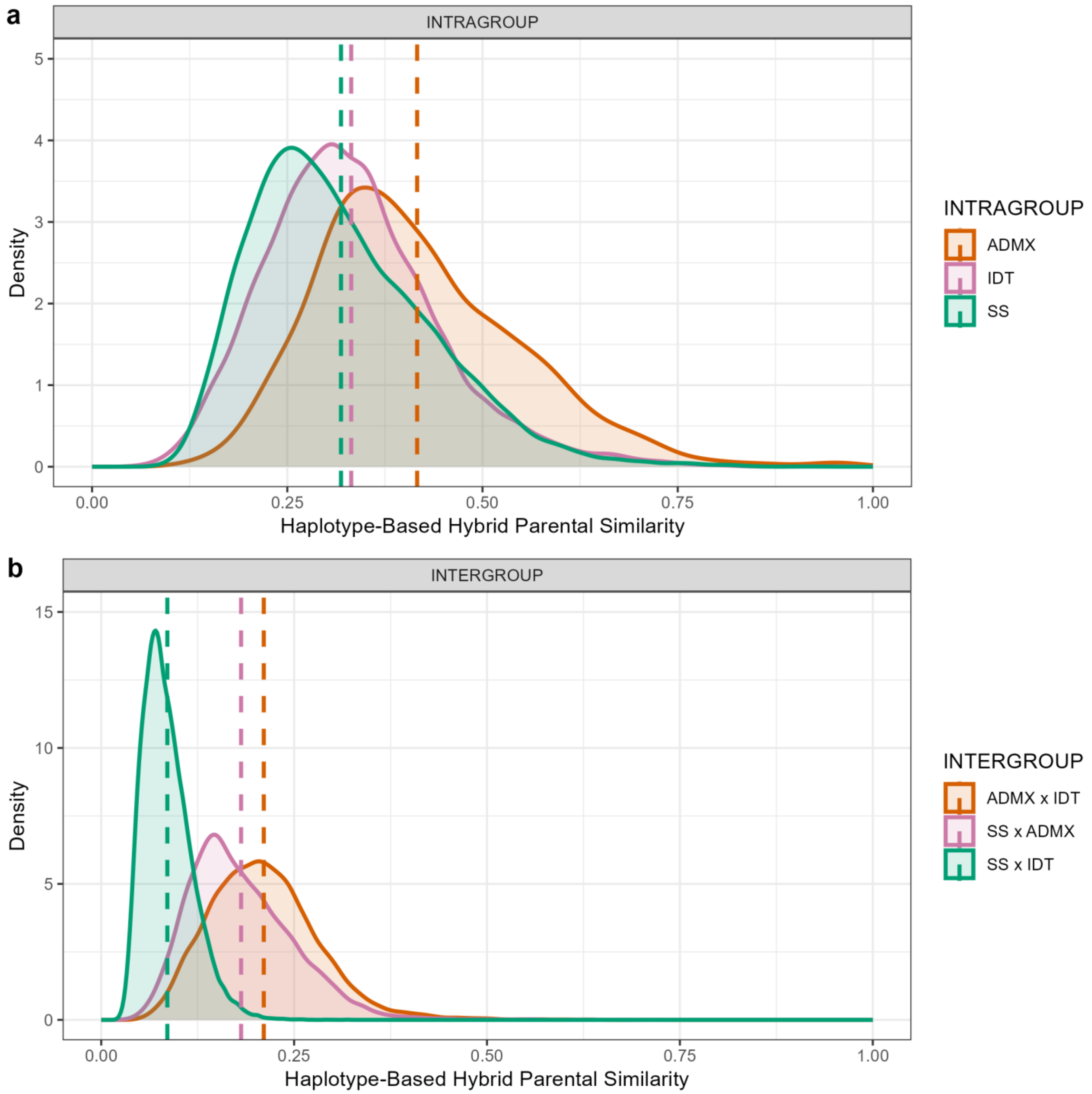

***Supplementary Fig. 7*** *Density diagrams of all pairwise combinations for the haplotype-based parental similarity method (RIB). Density plots are separated according to the cross type: intragroup (top) or intergroup (bottom). Vertical line represents the average of the specific category.*

90

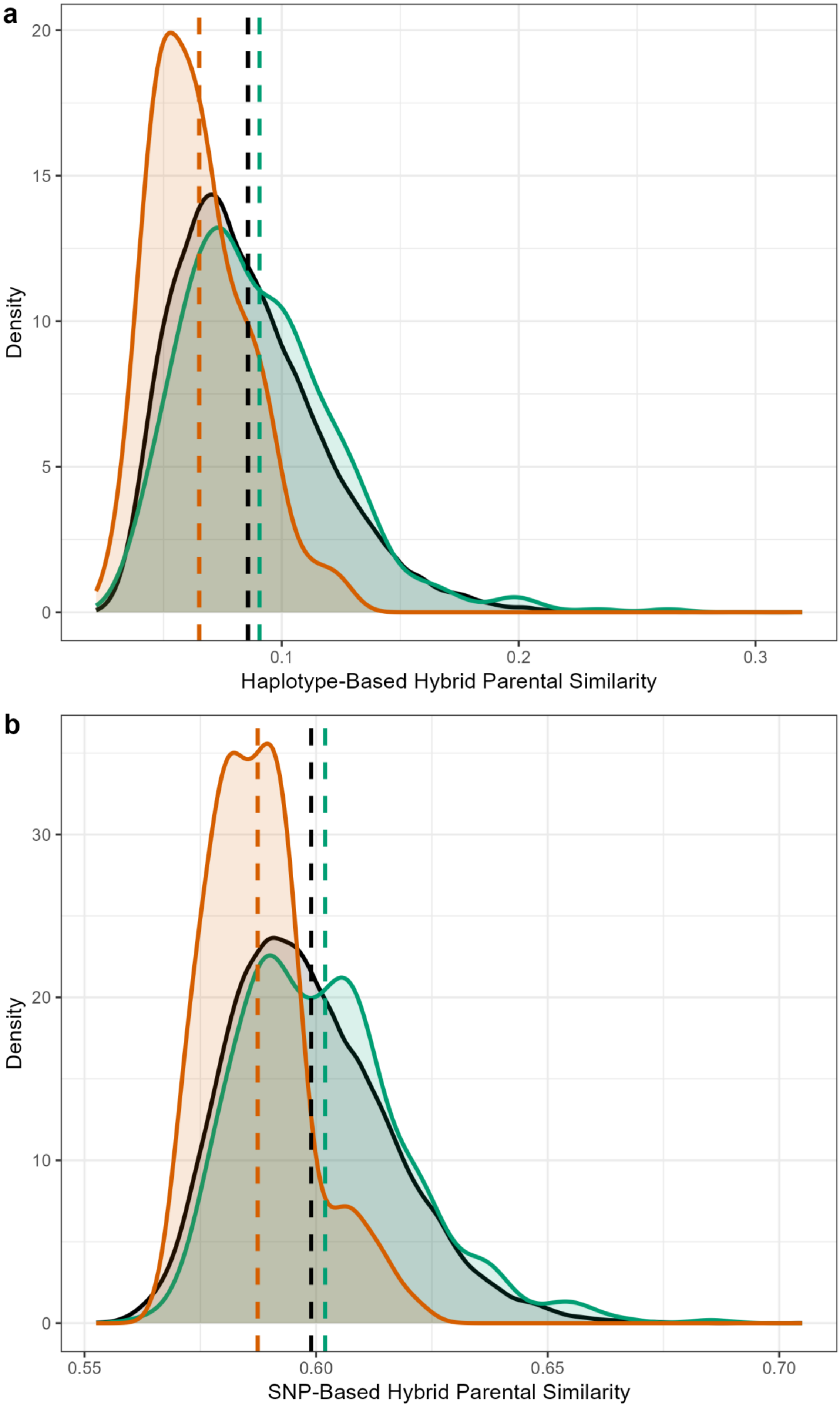

***Supplementary Fig. 8*** *Density diagrams of all pairwise Stiff Stalk x Iodent crosses for the haplotype-based parental similarity method (RIB) (top) and the SNP-based method (IBS) (bottom). Vertical line represents the average of the specific category. Color scheme separates the possible non-tested intergroup crosses (black), to the hybrids tested (green), and to the hybrids tested specifically in 2024 (orange).*
